# Sharp waves, bursts, and coherence: Activity in a songbird vocal circuit is influenced by behavioral state

**DOI:** 10.1101/2024.09.03.610933

**Authors:** Corinna Lorenz, Anindita Das, Eduarda Gervini Zampieri Centeno, Hamed Yeganegi, Robin Duvoisin, Roman Ursu, Aude Retailleau, Nicolas Giret, Arthur Leblois, Richard H. R. Hahnloser, Janie M. Ondracek

## Abstract

Similar to motor skill learning in mammals, vocal learning in songbirds requires a set of interconnected brain areas that make up an analogous basal ganglia-thalamocortical circuit known as the anterior forebrain pathway (AFP). Although neural activity in the AFP has been extensively investigated during awake singing, very little is known about its neural activity patterns during other behavioral states. Here, we used chronically implanted Neuropixels probes to investigate spontaneous neural activity in the AFP during natural sleep and awake periods in male zebra finches. We found that during sleep, neuron populations in the pallial region LMAN (lateral magnocellular nucleus of the nidopallium) spontaneously exhibited synchronized bursts that were characterized by a negative sharp deflection in the local field potential (LFP) and a transient increase in gamma power. LMAN population bursts occurred primarily during non-rapid eye movement (NREM) sleep and were highly reminiscent of sharp-wave ripple (SWR) activity observed in rodent hippocampus. We also examined the functional connectivity within the AFP by calculating the pairwise LFP coherence. As expected, delta and theta band coherence within LMAN and Area X was higher during sleep compared to awake periods. Contrary to our expectations, we did not observe strong coherence between LMAN and Area X during sleep, suggesting that the input from LMAN into Area X is spatially restricted. Overall, these results provide the first description of spontaneous neural dynamics within the AFP across behavioral states.

**Significance statement:** Although cortical and basal ganglia circuits are known to be required for learning in both mammals and birds, little is known about the ongoing spontaneous activity patterns within these circuits, or how they are modulated by behavioral state. Here we prove the first description of cortical-basal ganglia network activity during sleep and awake periods in birds. Within the pallial area LMAN, we observed population-wide bursting events that were highly reminiscent of hippocampal sharp-wave ripple (SWR) activity, suggesting that large-scale population events have diverse functions across vertebrates.

## Introduction

During vocal learning, a juvenile bird transitions from acoustically simple, highly variable “subsongs” to complex and stereotypical adult songs through a process of motor learning (Kollmorgen et al., 2020; Konishi, 1965, 1970; McCasland & Konishi, 1981). Critically involved in this learning process is a set of interconnected brain areas that make up a basal ganglia-thalamocortical circuit (Fig. 1A) known as the AFP (Andalman & Fee, 2009; Bottjer et al., 1984; Brainard & Doupe, 2000; Farries & Perkel, 2002; Scharff & Nottebohm, 1991)

**Figure 1:**
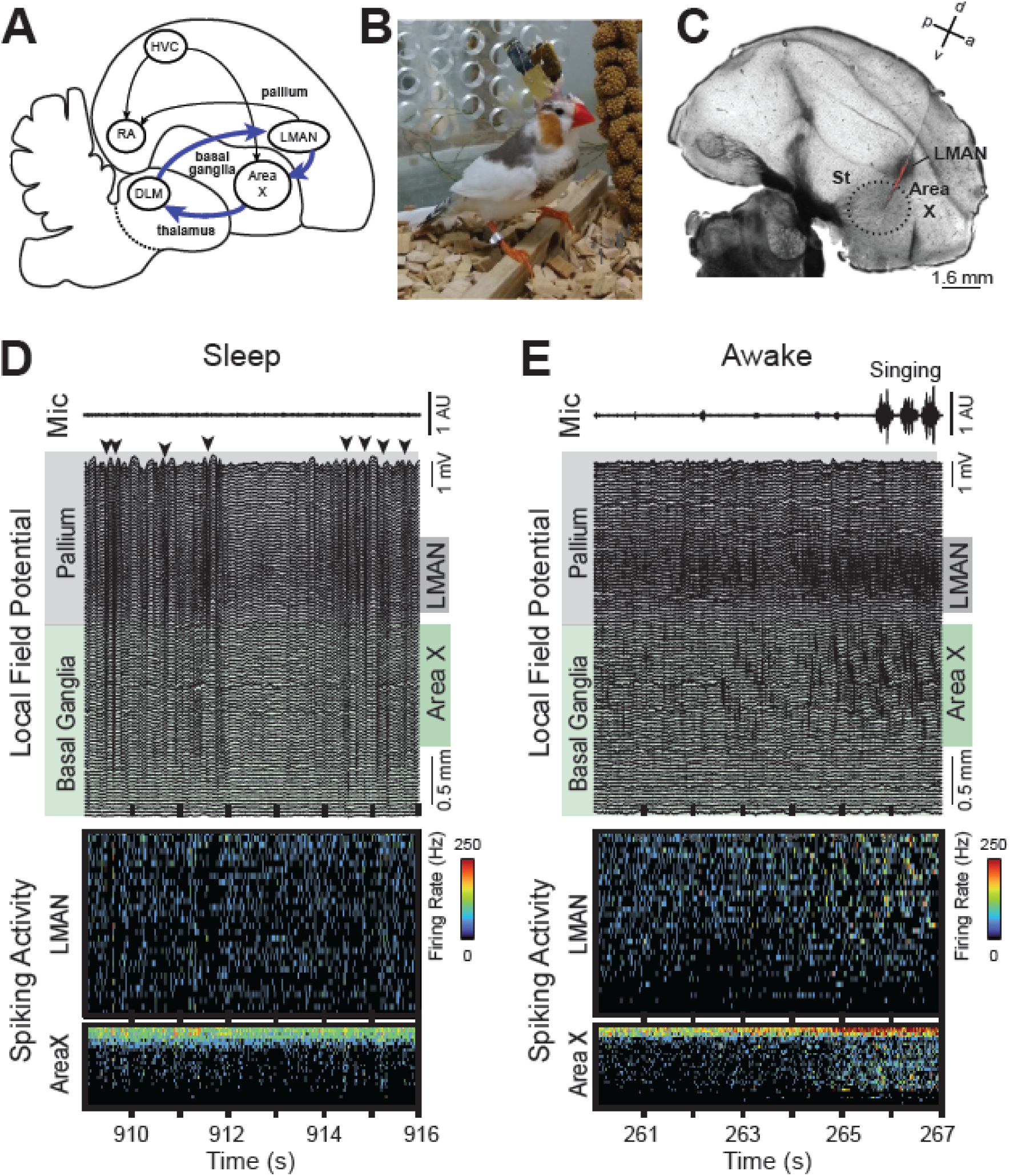
Pallial-basal ganglia dynamics during sleep and awake behavior in a songbird. **(A)** Schematic depicts the basal ganglia-thalamocortical circuit (blue arrows). Black arrows indicate other important connections of the song system. HVC, proper name; RA, robust nucleus of the arcopallium; LMAN, lateral magnocellular nucleus of the nidopallium; Area X, proper name; DLM, medial dorsolateral nucleus of the thalamus. **(B)** Image of a male zebra finch implanted with a Neuropixels1.0 probe. **(C)** Histological brain slice depicting the Neuropixels electrode track (red DiI fluorescence). Area X is indicated with a dotted black line. St, striatum; d, dorsal; a, anterior; v, ventral; p, posterior; for other abbreviations, see (A). Histology from ZF-2, cut at room temperature on a vibratome. **(D)** 7 s segment of the bandpass-filtered (0.5-250Hz) local field potentials (LFP) recorded during putative sleep in subadult ZF-1 and corresponding microphone signal (top, arbitrary units (AU)). Black arrowheads (LFP, top) indicate large amplitude negative LFP deflections. Recording sites were located in the pallium (gray shading) and basal ganglia (green shading) and are organized according to depth. Dark gray box (LMAN, right side) indicates LFPs that correspond to LMAN; dark green box (Area X, right side) indicates LFPs that correspond to Area X. Below, corresponding spiking activity for identified LMAN or Area X neurons. Firing rate is indicated with the color scale, and neurons are sorted according to firing rate. Note how neural activity transitions from sparse and synchronous firing aligned to the LFP troughs to asynchronous firing at 912-914 s. **(E)** 7 s segment of the bandpass-filtered LFP recorded during active daytime behavior in ZF-1. Figure conventions same as in (D). Note the calling and singing behavior recorded on the microphone signal (top) and the corresponding activity in LMAN and Area X. Note also the absence of large amplitude negative deflections in the LFP as observed during putative sleep.

Within the AFP, the pallial structure LMAN exerts a direct influence on vocal output through its projection to premotor area RA (robust nucleus of the arcopallium), presumably modulating downstream neural activity through bursts of spikes (Kao et al., 2008; Kojima et al., 2013; Palmer et al., 2021). Through this influence, LMAN drives behavioral variability that can be used for learning (Tumer & Brainard, 2007). Area X, part of the avian striatum (Person et al., 2008; Pfenning et al., 2014), is thought to receive an efference copy of activity in LMAN (Andalman & Fee, 2009; Vates et al., 1997; Vates & Nottebohm, 1995), thus placing Area X in a position to evaluate LMAN exploratory activity in the context of the ongoing song via a dopaminergic reward signal (Duffy et al., 2022; Kasdin et al., 2025; Qi et al., 2025).

While the AFP shares many similarities with the mammalian corticostriatal circuit, including cell types, neural connectivity, and gene expression patterns (Bolhuis et al., 2010; Gale & Perkel, 2010; Murugan et al., 2013), it is anatomically distinct from the mammalian circuit.

For example, the avian pallium (including LMAN) displays a widely different anatomical organization compared to mammalian cortex and is remarkable due to its lack of cellular layers. Similarly, the songbird Area X is an aggregate of neural populations that are typically segregated across various nuclei in mammalian basal ganglia (BG; (Person et al., 2008)). Despite these differences, the overall function of the circuit seems to be similar across birds and mammals, and many studies have shown that corticostriatal circuits are required for learning in both birds (Ali et al., 2013; Andalman & Fee, 2009; Bottjer et al., 1984; Olveczky et al., 2005; Scharff & Nottebohm, 1991; Tumer & Brainard, 2007; Zai et al., 2020) and mammals (Kawai et al., 2015; Lemke et al., 2021; Santos et al., 2015).

Though the anatomical similarities and differences between these circuits have been well characterized (Gale & Perkel, 2010), it remains unclear how these circuits compare in terms of neural activity. Spontaneous network activity that occurs during non-task-oriented behaviors, such as sleep or non-active awake periods, might be one way to broadly compare circuit dynamics across species. While aspects of the mammalian corticostriatal circuit have been well-studied in terms of neural activity across behavioral states (D. Liu et al., 2020) and sleep (Mizrahi-Kliger et al., 2018, 2020), including its role in the control of sleep and wakefulness (Lazarus et al., 2013) comparatively little is known about ongoing activity within the AFP. With the exception of one study that characterized the presence of high gamma oscillations in Area X during sleep (S. Y. Yanagihara & Hessler, 2012), spontaneous neural activity patterns in the AFP - and how they are modulated by behavioral state - remain largely unexplored.

We used a new approach (Lorenz, 2024) to chronically implant Neuropixels1.0 probes (Jun et al., 2017) in order to investigate neural activity in the pallial-striatal network of the AFP during periods of natural sleep and awake behavior in seven male zebra finches. By characterizing spontaneous neural spiking activity in relation to ongoing LFP oscillations in this avian pallial-striatal circuit, we gain insight into the different regimes of neural activity that are present within this circuit across different behavioral states, and which helps inform further studies of the analogous corticostriatal network in mammals.

## Materials and Methods

### Subjects

Seven male zebra finches (*Taeniopygia guttata*) were used in this study; data were pooled across two laboratories in France (the Neuroscience Paris-Saclay Institute and the Bordeaux Neurocampus). The age of the animals ranged from 76 days post-hatch (dph) to 232 dph at the time of recording. Where it was possible in the figures, we have identified the 3 youngest sub-adult animals (ZF-1, ZF-5, and ZF-7) with pink shading of their names. All animals were bred and housed in the animal facilities of the laboratories. During the experiment, animals were housed individually with water and food *ad libitum* under a 14:10 hr light/dark cycle.

Details about the individual animals can be found in Table 1.

**Table 1:** Information related to experimental animals. Age refers to the age at the recording time in days post hatch (dph). For all birds recorded in Bordeaux, data were recorded continuously throughout the night, and a 1-hour time period was selected for the analysis in this paper. The times indicated in the “Start of Recording” column indicates the start of the 1-hour recording time for those birds. * For some Paris birds, the dark period (light off) time was initiated earlier than normal and recording sessions started in the late afternoon (in the dark). LFP, local field potential SU, spiking unit (including single and multi-unit activity from high-pass filtered voltage traces) ** These animals died during the Covid laboratory lock-down in 2020 and it was not possible to retrieve the brains in time to perform histology. Electrode reconstruction was then based on response mapping of the brain areas that was performed during the probe implantation, as well as by visualizing the sorted spiking data for LMAN and Area X, which was distinguishable on the basis of spike amplitude and rate. ***Different spikesorting pipelines were setup in Paris and Bordeaux and largely reflected the different use of Matlab versus Python. When the data were first acquired in Paris, Kilosort 1.0 (KS 1.0) was the only version available; the data were sorted with Kilosort 1.0 and curated with Phy. Later, as more data were acquired in Bordeaux, the analysis pipelines there incorporated the newer version of Kilosort 2.5 (KS 2.5) and reflected E.G.C.Z.’s work with the Spikeinterface (SI) development team.

| Z<br>F | Animal<br>ID | Age<br>(dph) | Rec<br>location | Start of<br>recording | Data<br>available | # LMAN<br>units | # Area X<br>units | Histology | Spike-<br>sorting*** |
| --- | --- | --- | --- | --- | --- | --- | --- | --- | --- |
| 1 | g4r4 | 84 | Paris | 03:58<br>pm* | LFP+SU | 30 | 26 | Vibratome | KS 1.0<br>+Phy |
| 2 | r15v15 | 125 | Paris | 8:42 pm | LFP+SU | 32 | 6 | Vibratome | KS 1.0 +<br>Phy |
| 3 | j8v8 | 124 | Paris | 05:18 am | LFP+SU | 40 | 19 | Missing** | KS 1.0 +<br>Phy |
| 4 | Redred | 177 | Bordeaux | 05:30 am | LFP+SU | 28 | 7 | Cryostat | KS 2.5 +<br>SI |
| 5 | r5n5 | 85 | Paris | 5:21 pm* | LFP+SU | 6 | 3 | Vibratome | KS 1.0 +<br>Phy |
| 6 | black/cyan | 232 | Bordeaux | 04:17 am | LFP | - | - | Cryostat | - |
| 7 | no_label | 76 | Bordeaux | 00:28 am | LFP | - | - | Missing** | - |

All experimental procedures were approved by the French Ministry of Research and the ethical committee “Paris-Sud et Centre’’ (CEEA No. 59, project 2017-25) and “Poissons Oiseaux Nouvelle Aquitaine” (CEEA No. 73, project S73) and performed under the license 2015-25 and APAFIS#13413-2018020713145795.

### Surgery and experiments

#### Chronic Neuropixels implantation surgery

The day of the surgery, food and water sources were removed from the cage approximately 30 minutes before the start of the surgery. Animals were anesthetized with isoflurane (0.6-1.5% inhalation) and placed in the stereotactic apparatus once the toe flexion reflex was no longer observed. Feathers were removed from the scalp and the area was disinfected with ethanol and betadine. 0.05 microliters of diluted lidocaine (0.5mg/kg) was administered subcutaneously and a local anesthetic (0.5 g Emla® creme) was applied to the exposed skin. The scalp was resected and the exposed skull was prepared for the implant by drilling several small holes in the outer bone layer. A small craniotomy contralateral to the final implant was made to place a silver wire in contact with the dura that served as the electrical ground.

The main craniotomy was made on the right hemisphere at approximately 1.7 mm lateral to the superior sagittal sinus and 4.5 mm anterior to lambda, the confluence of sinuses, with the flat part of the anterior skull rotated to a 50-degree angle. The positions of LMAN and Area X were mapped with high-impedance single-electrodes in order to identify the brain areas based on the firing rate properties thereof.

A Neuropixels probe version 1.0 (Jun et al., 2017) was implanted 4 to 5 mm deep to penetrate the estimated center of LMAN and Area X. The custom-made holder that carried the Neuropixels probe was fixed to the skull with dental cement and the craniotomy was sealed with a two-part silicone gel (Dow Dowsil^TM^, PN 3-4680) before it was covered with a custom-made casing. In order to fix the Neuropixels headstage to the bird’s head in a natural position, the bird’s head was rotated to a 35-degree angle in the stereotaxic during the fixation. Finally, the ground wire was connected and open holes along the implant were sealed with fast drying two-component silicone gel (Kwik-Sil).

The weight of the implant (∼1.9 g) was immediately offset by a weight-relieving system. The headstage was connected to an external counterweight using a nylon thread that was routed through simple guide points positioned above and to the side of the setup, forming an inverted U-shape. This allowed the counterweight to remain outside the recording area while still providing vertical support and offsetting the implant’s weight. The counterweight was manually adjusted to ensure that the animals could move comfortably and were not burdened by the implant.

#### Histology

Post-hoc control of the electrode location in the brain was performed through histological examination of brain tissue. At the end of the experiment, birds were given a lethal injection (i.p) of Exagon (pentobarbital sodium, 400 mg/mL), and in some cases perfused intracardially (0.01 M Phosphate-Buffered Saline (PBS) followed by 4% paraformaldehyde (PFA)). The brain was removed and post-fixed in 4% PFA for at least 24 hr. For 2 birds recorded in Bordeaux, the brains were subsequently cryoprotected in 30% sucrose and cut using a freezing microtome; for 3 birds recorded in Paris, brains were cut fresh at room temperature using a vibratome. In all cases, the right hemisphere was cut in the sagittal plane at a thickness of 50-80-microns. Images of the individual brain slices were taken using bright-field and epiflurescence microscopy and analyzed using ImageJ software (Rasband WS, NIH, Bethesda, Maryland, USA). The electrode tract, as indicated by colored/fluorescent dye (Dil fluorescent dye; DilC_18_(3)), was clearly visible on consecutive brain slices (see Fig. 1C; tissue from ZF2 and prepared on a vibratome).

2 birds died during the Covid-19 laboratory lockdown, and we were not able to retrieve the brains in time to perform histology (Table 1). For these animals, the channel assignment was performed on the basis of the electrophysiology, as described in detail below.

#### Electrode channel mapping

We defined the boundaries of LMAN and Area X based on darkness of the tissue in regular light or the fluorescent background, as these two brain areas showed a clear contrast with surrounding tissue due to higher myelination, cell density, and size (Nixdorf-Bergweiler et al., 2007). The reconstruction of electrode channels to be assigned to LMAN or X was performed based on the location of the electrode tip on the histological pictures, the coordinates of the channels on the electrode and the above-mentioned boundaries of LMAN and X. The limits of LMAN were also confirmed based on spontaneous activity, including more bursts of large spikes in LMAN as compared to surrounding tissue.

For the 2 birds with missing histology, we relied heavily on these electrophysiological properties, including the spike amplitude, rate, and response mapping that was performed during the surgical probe implantation. Generally, LMAN channels were associated with larger spike amplitudes, whereas Area X channels had higher spike rates.

#### Experimental design

Animals were housed individually in sound-attenuated recording chambers and neural recordings were performed in sessions (one to 24 hours of continuous neural recording).

#### Electrophysiology

Extracellular activity was recorded with the chronically implanted Neuropixels 1.0 probe which splits the signal on chip and samples the spike signal (bandpass filtered at 0.3 and 10 kHz) at 30 kHz and the LFP signal (bandpass filtered at 0.5 and 500 Hz) at 2500 Hz. Audio signals were simultaneously recorded using a wall-attached microphone, pre-amplified and digitized at 10, 20 or 32 kHz depending on the bird.

#### Sleep Recordings

Sleep-related activity was recorded either continuously throughout the night (n=3 birds; Table 1) or at the beginning or end of the night (n=4). In the case of continuous recordings, a one-hour segment was selected by visually inspecting the microphone data for a consolidated quiet period indicative of sleep. In 5 out of 7 birds, sleep-related activity was recorded with the light off during the normal dark period; however, in 2 birds (see Table 1), the dark period was initiated earlier than normal, and recording sessions started in the late afternoon (with light off).

#### Awake Recordings

For 4 out of the 7 birds (ZF-1, ZF-2, ZF-3 and ZF-5), we also examined brain activity that occurred during normal daytime hours in the light. The behavior of the birds during this time was characterized by singing, active movement, and periods of rest. Because it was possible that birds were napping during periods of daytime rest, we classified the awake data into two behavioral states on the basis of sounds detected from the microphone. “Light on active” periods corresponded to microphone data which contained singing or sounds related to movements. (These periods were distinct from movement-related voltage artifacts which were detected as described below in section “Sound detection and movement artifact identification”). “Light on non-active” periods corresponded to microphone data where no sounds were detected. These could be periods where the bird was awake but not moving or in a state of quiet daytime rest (see section “Sound detection and movement artifact identification” and Table 2 for more information).

**Table 2:** Average coherence values for all birds (n = 7) used for analysis in. **Figure 4 C.** Frequency ranges: Delta (1 – 4 Hz), Theta (4 – 12 Hz), Beta (12 – 30 Hz), Low gamma (30 – 90 Hz), High gamma (90 – 140 Hz). N corresponds to the number of LFP pairs.

| Brain Area | Frequency range | Median coherence across channels during SWS (Min, Max values) | Median coherence across channels during REM (Min, Max values) | Wilcoxon signed rank test |  |
| --- | --- | --- | --- | --- | --- |
|  |  |  |  | z value | p value |
| **LMAN (N=147)** | **Delta** | 0.965 (0.861, 0.996) | 0.872 (0.633, 0.989) | 10.52 | 7.15E-26 |
|  | **Theta** | 0.968 (0.876, 0.995) | 0.893 (0.754, 0.988) | 10.52 | 7.15E-26 |
|  | **Beta** | 0.821 (0.521, 0.969) | 0.680 (0.311, 0.963) | 10.52 | 7.15E-26 |
|  | **Low gamma** | 0.500 (0.119, 0.854) | 0.484 (0.106, 0.862) | 6.244 | 4.26E-10 |
|  | **High gamma** | 0.397 (0.074, 0.769) | 0.451 (0.063, 0.806) | -5.67 | 1.43E-8 |
| **Area X (N=2275)** | **Delta** | 0.606 (0.043, 0.946) | 0.409 (0.006, 0.888) | 39.34 | 0.00E+00 |
|  | **Theta** | 0.671 (0.131, 0.977) | 0.561 (0.101, 0.939) | 41.00 | 0.00E+00 |
|  | **Beta** | 0.394 (0.062, 0.928) | 0.304 (0.046, 0.912) | 41.28 | 0.00E+00 |
|  | **Low gamma** | 0.189 (0.024, 0.770) | 0.178 (0.023, 0.703) | 31.30 | 5.44E-215 |
|  | **High gamma** | 0.146 (0.021, 0.691) | 0.139 (0.020, 0.648) | 24.77 | 1.69E-135 |
| **LMAN – Area X (N=1274)** | **Delta** | 0.169 (0.003, 0.820) | 0.163 (0.011, 0.782) | 10.50 | 9.05E-26 |
|  | **Theta** | 0.244 (0.011, 0.818) | 0.234 (0.008, 0.749) | 4.79 | 1.70E-6 |
|  | **Beta** | 0.107 (0.013, 0.464) | 0.117 (0.015, 0.480) | -8.37 | 5.71E-17 |
|  | **Low gamma** | 0.069 (0.019, 0.153) | 0.077 (0.020, 0.172) | -14.35 | 1.06E-46 |
|  | **High gamma** | 0.055 (0.014, 0.171) | 0.040 (0.010, 0.139) | 30.84 | 8.16E0-209 |

Note that by segmenting activity solely based on microphone-detected sounds, we may neglect neural activity changes occurring in Area X and LMAN prior to song initiation (Hessler & Doupe, 1999a; Woolley et al., 2014),. Such premotor activity could therefore be misclassified as occurring during “Light on non-active.” However, this misclassification is likely uncommon, as zebra finches are generally behaviorally active in the periods surrounding song bouts (Zann, 1996).

### Statistical analysis

#### Data preprocessing and spike clustering

For neural spiking analysis, spike sorting was performed in four of the seven birds on all active channels using Kilosort 1.0 or Kilosort 2.5 (Pachitariu et al., 2016; see Table 1 for more information) and manually curated using phy (https://github.com/kwikteam/phy) in four birds (ZF-1, ZF-2, ZF-3 and ZF-5) or automatically curated using SpikeInterface (script available in our data repository) in one bird (ZF-4). Regardless of the sorting process, a user-based quality check of the final dataset ensured that the quality of sorted units was satisfactory and homogeneous.

#### Sound detection and movement artifact identification

We identified periods of putative awakening during the light-off period by analyzing the microphone recordings. We calculated the root mean square (RMS) of the microphone signal and manually defined a threshold that was just above the noise level (Fig. 3A, Fig. S1). Each threshold crossing identified a putative movement-related sound, and 1-s bins during which sound was detected were excluded from further analysis.

**Figure 2:**
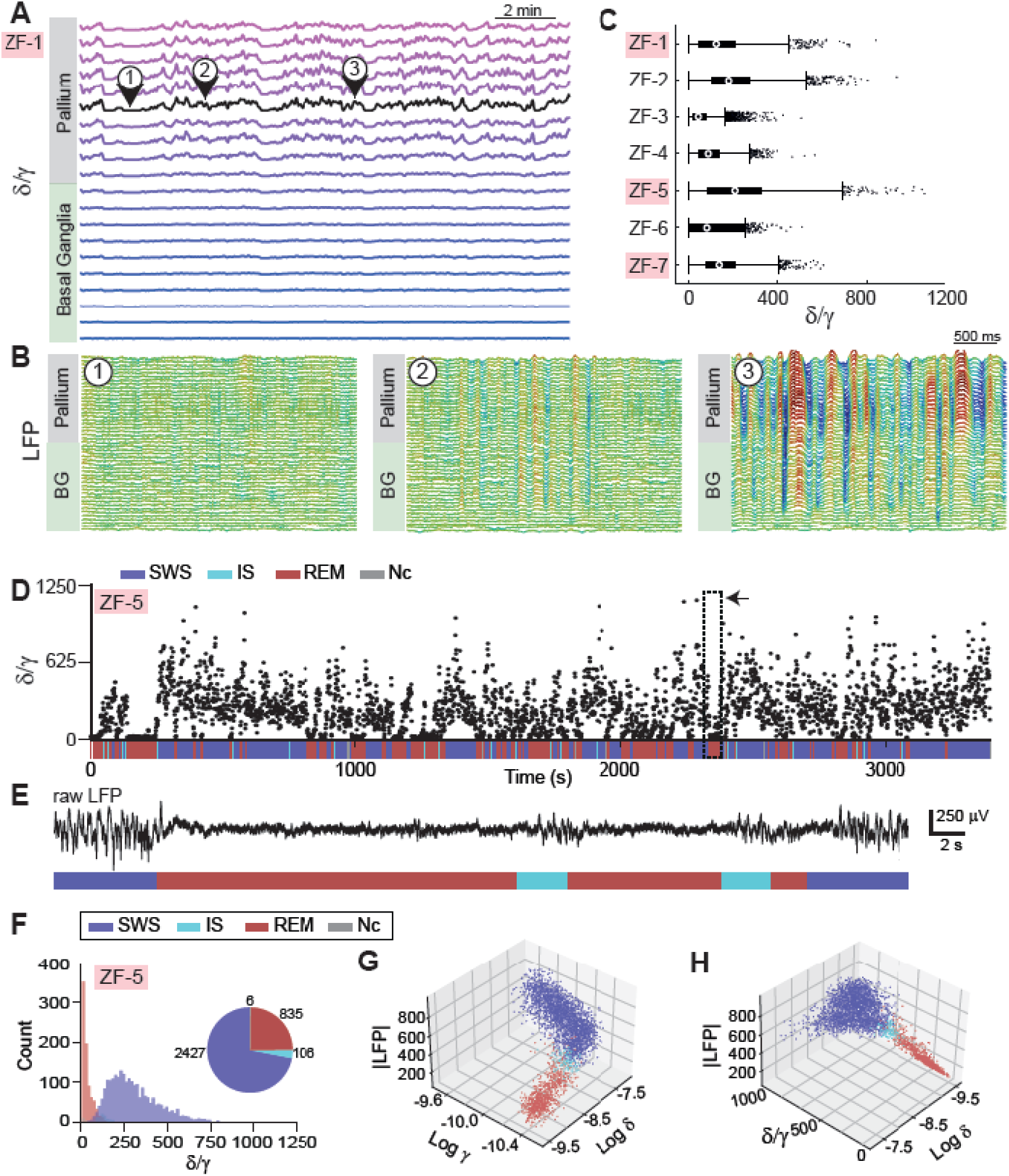
Identification of sleep stages and representative neural dynamics. (**A)** Ratio of δ/γ power for consecutive depths along the Neuropixels shank. Each trace is separated by 200 um. Black trace indicates the δ/γ trace calculated for a centrally located pallial (LMAN) LFP. Numbers and arrows indicate low (1), middle (2), and high (3) δ/γ values. δ/γ values calculated for a 3-s LFP window, as depicted in (B). Data from sub-adult ZF-1 (pink shading). **(B)** 3-second segments of local field potential (LFP) activity from pallial and BG sites for the low, middle, and high δ/γ values indicated in (A). Each trace is separated by 80 um. Note the large amplitude slow wave dynamics present for high δ/γ values indicated in (3). Linear color scale for visualization purposes only. Data from sub-adult ZF-1. **(C)** Box plots indicate average δ/γ values calculated in 3-s windows from a centrally located LMAN channel for the duration of the sleep phase per bird. The left and right edges of the box represent the 25th and 75th percentile, respectively, and the middle dot represents the median. Whiskers extend to the most extreme datapoints not considered outliers, and datapoints beyond this range are indicated as dots. Sub-adult animals are indicated with pink shading of their names. **(D)** δ/γ values for an entire recording duration for an example bird (sub-adult ZF-5) plotted with the identified sleep stages as colored bars below. SWS, slow wave sleep; IS, intermediate sleep; REM, rapid eye-movement sleep; Nc, not categorized. Dotted black box indicated with the black arrow is enlarged in (E). **(E)** 50 s trace of raw LFP from the LMAN channel used for sleep staging (see Methods for details). Color-coded bars indicate classified sleep stages. Note how SWS segments correspond to a large amplitude LFP signal and REM segments correspond to a low amplitude LFP signal. **(F)** Histogram of δ/γ values grouped according to identified sleep states together with the proportions of sleep stages depicted in a pie chart for the bird in (D). Note how SWS segments correspond to high δ/γ values and REM segments correspond to low δ/γ values. **(G, H)** Plots show that sleep state clusters are segregated in 3-D space according to the features used for classification, e.g., mean LFP magnitude, the delta (δ) and gamma (γ) power, δ/γ values. Each dot represents a 1-s snippet of sleeping data (resolution of the automated sleep staging procedure). Same bird as in (D-F).

**Figure 3:**
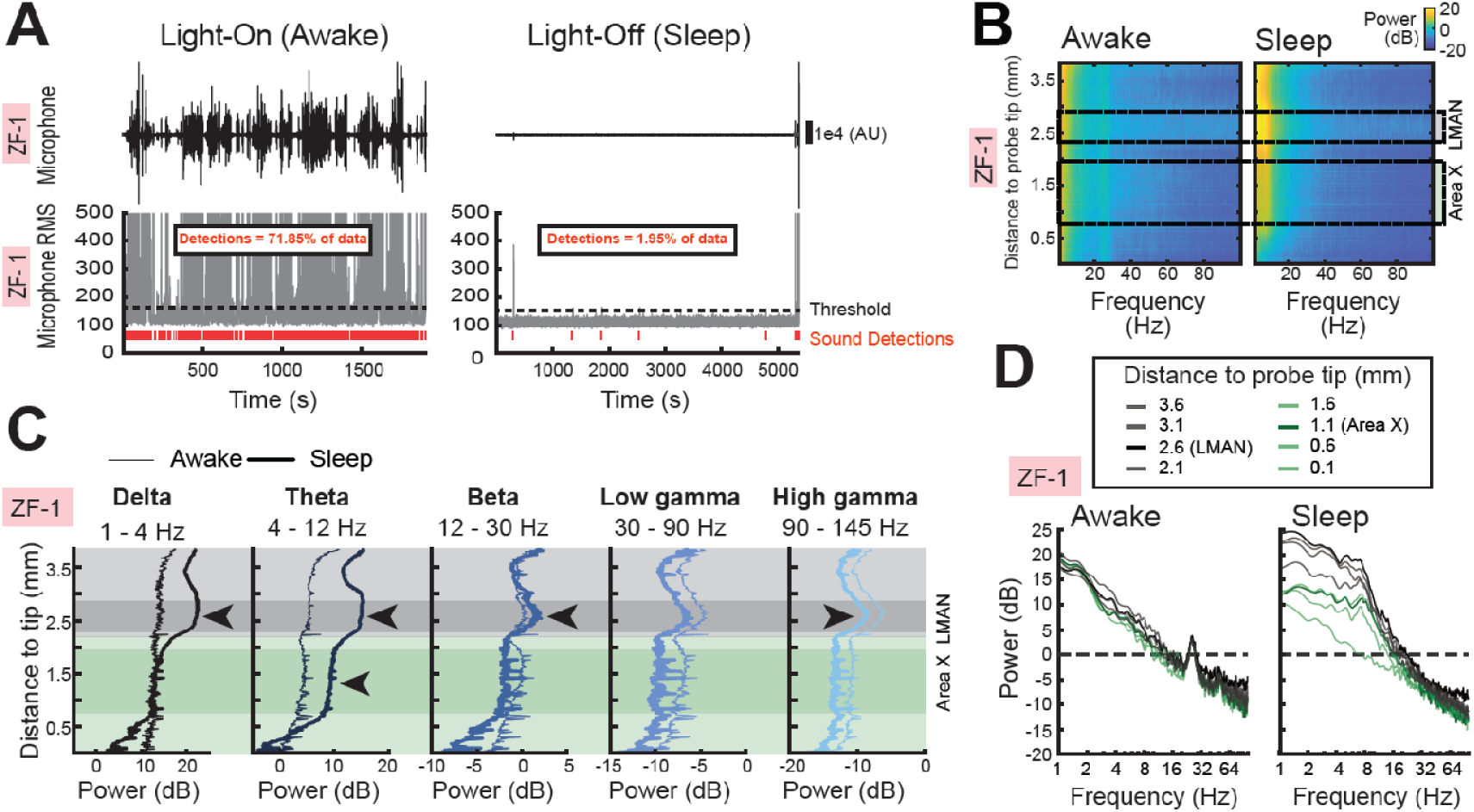
Sleep-related modulation of low-frequency power is higher in pallium compared to the striatum. (**A**) Comparison of microphone signals recorded during the light on (awake) period and the light off (sleep) period. Top traces (black) indicate the raw microphone signal; scale is the same across recordings (arbitrary units, AU). Bottom traces (gray) indicate the root mean square (RMS) of the microphone signal. Black dotted line indicates the manually set threshold, and threshold crossings indicate sound detections (red lines). The overall percent of data for which sounds were detected is stated (top). Recording from sub-adult ZF-1. Microphone data from other birds can be found in Fig. S1. **(B)** The color-coded power spectrum density (PSD) of the LFP signal for each electrode is displayed, organized by depth (dorsal to ventral), for both the awake state (left) and sleep state (right) for sub-adult ZF-1. Warm color indicates high power; cool colors indicate low power. The electrode depths corresponding to LMAN (gray box, dashed line) and Area X (green box, dashed line) are indicated. The analysis is based on the average PSD derived from 10 randomly selected 10-sec windows. **(C)** Power distributions across electrode sites provide a detailed comparison between awake (thin lines) and sleep (thick lines) for different frequency bands. LMAN exhibits higher gamma band power (90-145 Hz) during awake states compared to sleep states (LMAN; black arrowheads). In contrast LMAN exhibits higher power in low frequency bands (1-30 Hz) during sleep compared to awake states (LMAN; black arrowheads). A difference in sleep and awake states in Area X is largely absent, with the exception of the theta band power (4-12 Hz), which showed a slight increase in two birds during sleep (Area X; black arrowhead). **(D)** The marginal distribution of power across depths is displayed for awake and sleep states for each bird. Note that the distances in the legend refer to the distance from the probe tip, (i.e., 3.6 mm from the probe tip). The black line indicates depth associated with LMAN (i.e., 2.6 mm); note how it is strongly modulated by sleep. Gray lines indicate other depths associated with the pallium. The dark green line indicates the depth associated with Area X; light green lines indicate other depths associated with the BG. Dotted line indicates 0 dB. Spectral data for other birds are depicted in Fig. S4.

During the light-off recordings, it is possible that the bird could awaken briefly but not move, which would not be identified as a putative awakening by our sound detection approach.

Although we are not able to quantify the extent to which such events may contribute to our dataset, other papers which used video recordings to confirm sleep states have reported discarding between 2% (Yeganegi & Ondracek, 2025) and 3% (Yeganegi & Ondracek, 2023) of sleep EEG data due to movement-related artifacts. Across our birds, the percentage of data discarded using the sound detection method ranged from 0.12% to 6.02% (see Fig. S1 for exact values). Therefore, although our approach using sound detections to identify putative awakenings cannot identify quiet awakenings in the dark, we estimate that this would affect at most an additional 3% of the data. We evaluate and discuss the potential inclusion of quiet awakenings in our sleep analysis in the Discussion and in Text S1.

For the analysis of awake neural activity, we similarly leveraged the microphone recording to distinguish between silent periods and periods of activity. Periods where sounds were detected were identified as “Light on active” periods, and silent periods were identified as “Light on non-active.”

The awake data often exhibited movement-related artifacts in the LFP which we did not observe in the putative sleep recordings. These daytime movement artifacts were characterized by synchronous fluctuations across all channels. To address this, we applied a modified version of the Artifact Subspace Reconstruction (ASR, (Chang et al., 2020; Kothe & Makeig, 2013) algorithm to identify and exclude artifact-contaminated periods. Briefly, we performed an eigenvalue decomposition of the data covariance matrix. Components associated with large eigenvalues indicative of high-variance noise, were flagged as potential artifacts. These eigenvalues were identified using a threshold defined as a multiple *q* of the median eigenvalue. To detect specific times of contamination, we reconstructed the LFPs using only the artifact-related components and computed the summed signal across all channels. Time bins where this signal exceeded a threshold—defined as the mean plus *k* times the standard deviation—were marked as contaminated and excluded from further analysis. Both *k* (ranging between 3 and 12) and *q* (ranging between 10 and 30) were manually tuned for each bird to conservatively exclude movement artifacts without discarding valid neural activity.

#### Sleep staging

The sleep staging was performed using an unsupervised k-means clustering algorithm previously developed in songbirds to classify sleep data on the basis of multiple EEG features (Canavan & Margoliash, 2020; Low et al., 2008). This algorithm classifies sleep into one of 3 states: SWS, REM sleep, and intermediate sleep (IS) which has been compared to the mammalian NREM sleep stage N2 and often contains a mixture of low-amplitude delta and higher-frequency elements such as theta (Canavan & Margoliash, 2020). Our use of this method diverged from its original implementation in the following ways: 1) we used LFP instead of EEG to compute the sleep stages, 2) our exclusion of awake periods was based on sound detection from the microphone (as described above), and did not include manual scoring of video data to confirm that animals in the dark were sleeping prior to sleep state classification, as was done in (Canavan & Margoliash, 2020). The LMAN LFP channel used in the analysis was a centrally located LMAN channel along the dorsoventral axis of recording. Spectral parameters were computed from the multi-taper spectral analysis (Prerau et al., 2017) of the filtered (0.5 - 60 Hz) and down sampled (250 Hz) LFP using 3 s bins (1 s overlap). A small number of bins were not classified as REM, SWS or IS using the two-step clustering algorithm (Canavan & Margoliash, 2020) and were labeled ‘Not classified/Nc’. Nc classifications arise when a bin is classified as SWS in the first clustering round and then as REM in the second clustering round.

Our use of a centrally located LMAN LFP channel to classify sleep stages (instead of a superficially located EEG electrode) means that our sleep staging results may diverge slightly from values reported elsewhere (Canavan & Margoliash, 2020; Low et al., 2008; Yeganegi & Ondracek, 2023, 2025). However, LFP signals were filtered in the same way that EEG signals were filtered (i.e., between 0.5 and 60 Hz), which would eliminate high-frequency characteristics that might be present in an LFP recording but not in an EEG recording.

Another concern was that the LFP recording centered in LMAN might not reflect “global” sleeping activity. Indeed, recent work has shown that sleeping brain dynamics are not as global as previously thought, and different sleep states might occur in different brain regions, as was also reported for songbirds (Yeganegi & Ondracek, 2025). To investigate this in our own data (Fig. S2), we examined the differences in sleep classifications in a superficial LFP located in the hyperpallium (comparable to a superficially implanted EEG electrode) and compared it to a centrally located LMAN channel (Fig. S2A). We found that almost 90% of the sleep scoring matched between the LMAN channel and the superficial LFP channel (Fig. S2B), suggesting that the use of an LMAN channel to assess sleep state is a fair assessment of the ongoing sleep states that would have been recorded using a superficial EEG electrode. Of the 11% of mis-matched sleep states, the largest category involved a mismatch between SWS and IS, such that sleep states that were identified as SWS in LMAN were identified as IS in the hyperpallium (Fig. S2C).

### LFP Coherence Analysis

#### Data inclusion

For the coherence analysis we only used bins identified as either REM or SWS. Time points at which movements were detected, as described in the previous section (Sound detection and movement artifact identification) were excluded for the coherence analysis.

#### LFP pre-processing

First, we extracted the LFP data from *.bin* files utilizing SpikeInterface (https://spikeinterface.readthedocs.io/en/latest/overview.html) with phase shift correction using *spikeinterface.preprocessing.phase_shift*. Next, we removed artifacts from the LFP signal by establishing a threshold based on 80% of the maximum absolute trace value and interpolating the signal around the artifact time point with a margin of 20 samples before and after. Next, we z-scored the trace using *elephant.signal_processing.zscore*.

#### Coherence Analysis – Sleep recordings

Pairwise LFP coherence was computed from the z-scored LFP traces. We included every 4^th^ LFP channel (corresponding to a dorso-ventral linear distance of approximately 40 microns between them) spanning Area X and LMAN to reduce the computation time while still retaining enough spatial resolution to obtain a clear estimate of the spatial variation. We performed pairwise LFP coherence per identified state. For sleep recordings, this involved computing coherence for the SWS and REM states respectively. Specifically, we computed the magnitude squared coherence *c_1,2_* of LFP signals L1 and L2 as follows:

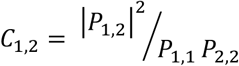

where *P_1,1_* and *P_2,2_* are power spectral density estimates of L1 and L2 using Welch’s method, and *IP_1,2_I* is the absolute value of the cross spectral density estimate. We used the *scipy.signal.coherence (*https://docs.scipy.org/doc/).

The accuracy of coherence magnitude estimates depends on 1) the length of the segments used for computation that is usually determined by the lowest frequency of interest (FOI) and 2) the total number of such segments obtained (equal to the total length of the signal being analyzed divided by the segment length). In our case, these parameters were constrained by our lowest FOI which was 1 Hz and the total duration of REM or SWS states in our recordings per bird. We selected a segment length of 3 sec for our analysis, since 3 sec segments were sufficient to give an accurate coherence estimate for our lowest FOI (1 Hz) and our total length of recordings corresponding to SWS or REM were large enough to obtain sufficient number of 3-sec segments.

After accounting for removal of movement artifacts and the analysis of only the REM and SWS segments, the total percentage of data analyzed in the coherence analysis per bird were: ZF1: 78.2%, ZF2: 78.4%, ZF3: 65.9%, ZF4: 76.2%, ZF5: 85.8%, ZF6: 77.4%, ZF7: 71.4%. Because the number of SWS segments were more numerous that the number of REM segments, for each channel comparison, we initially performed the coherence computation using a randomly selected number of SWS segments equal to the total number of REM segments. We then averaged across all SWS segments after ascertaining that the standard error of mean for the averaged coherence for all the SWS segments was low (< 0.05 for all channel comparisons for all birds) indicating low variation between randomly selected SWS segments.

In order to evaluate statistical significance in the coherency analysis between SWS and REM states, we first selected a minimum number of channels to compare that was the same for all birds for each brain area (LMAN = 7; equal to ∼240 microns, Area X = 26; equal to ∼1000 microns). This was necessary as the probe implantation was variable across animals and resulted in different number of total channel counts in the brain areas of different birds. Then we identified the most central channel in each brain area and included neighboring channels along the probe length. After pooling these channel comparisons across birds for SWS and REM states, we identified significant differences across brain states in the coherence analysis using a Wilcoxon Signed Rank test.

#### Coherence Analysis – Awake recordings

A similar procedure was followed for analysis of the awake daytime recordings (obtained from 4 out of the 7 birds). Here our goal was to determine whether behavioral state affected the coherence between brain areas. That is, we compared sleep-related activity to periods of Light on active and Light on non-active states. Only 3 out of the 4 birds (ZF-1, ZF-2, and ZF-4) had sufficiently long periods of both Light on active states to allow us to compare the pair-wise coherence analysis (see Table S1 for more information). Coherence values were pooled across birds as described above. To compare the awake and putative coherence results for these 3 birds, we performed statistical testing using a Friedman Test followed by the Nemeyni posthoc tests to account for multiple comparisons between non-independent samples (same channels and 4 different behavioral states).

#### Analysis of spiking variability

To assess how spike activity varied over time within LMAN and Area X during putative sleep, we computed the coefficient of variation (CV) in consecutive, non-overlapping segments of 10 seconds. Within each segment, we counted the total number of spikes (*ns*, from all units either in LMAN or in Area X) in sub-windows of 20 milliseconds. The CV was then computed as the standard deviation of spike counts divided by the mean spike count across all sub-windows within that segment (CV =*a*_ns_/*µ*_ns_). Because we were interested in the relationship between the temporal variability of spike activity within a population and how it related to putative sleep, we computed the δ/γ ratios for the LFP signal of the central electrode in LMAN in the same 10-second segment. We then used Pearson’s correlation coefficient (Pearson, 1895) and type-II linear regression (Ricker, 1973) to assess the relationship between δ/γ ratios and the CV of spike activation. The average firing rate of neurons in LMAN and Area X was calculated by determining the total spike count per second within the same windows where CV was computed. We performed this analysis for sleep recordings in five birds (ZF-1 to ZF-5), excluding segments that were contaminated by potential movement (see *Sound detection and movement artifact identification*).

#### Population synchronized burst detection and spectral analysis of associated LFP

For each neural unit (single or multiunit), bursts were identified when the Instantaneous Firing Rate (IFR) exceeded 100 Hz. The fraction of simultaneously bursting LMAN units was determined in a sliding 50 ms window with 10 ms step size. The resulting trace was smoothed using a Gaussian kernel with 15 ms standard deviation, and candidate events of synchronized bursts were classified as peaks exceeding the 99^th^ percentile.

We performed this analysis for sleep-related recordings of five birds (ZF-1 to ZF-5). Events that occurred during the 1-sec segments that were flagged as putative awakenings or Nc sleep states were excluded. Similarly, in daytime recordings, events that overlapped with LFP movement artifacts were also excluded (see *Sound detection and movement artifact identification*).

To create control events, we used random time points in the recording. For assessing the LFP around synchronized bursts and control time points, we selected the central LMAN LFP and compared raw traces that were offset-corrected by median subtraction. Interested in a high-resolution description of the non-stationary spectro-temporal content of the associated LFP, we applied a complex wavelet transform using complex Morlet (Gabor) wavelets (*cwt,* MATLAB, The MathWorks, Natick, MA). Scalograms were computed over a 1-sec window centered on each event. Vice versa, to assess the oscillatory context of these events, we calculated the power estimate in the δ- (1-4 Hz) and γ-band (30 – 90 Hz) in the 3-sec window around the event, the same bin size used for sleep staging, using Thomson’s multitaper power spectral density method (*pmtm*, MATLAB, The MathWorks, Natick, MA). For the high γ-band (90 – 145 Hz), we used a narrower window of 300 ms centered to the event.

#### Data and software availability

All research data as well as scripts to replicate the results are available in the corresponding repository: https://gin.g-node.org/E4-Leblois/npx_sleep_2024

Spectral analysis and visualizations in Fig. 3B-D and S4 were computed with an adapted code of the following repository https://github.com/cortex-lab/spikes.

## Results

### Distinct regimes of activity are present during avian sleep

We investigated neural activity in the pallium and basal ganglia of male zebra finches (n = 7) chronically implanted with Neuropixels 1.0 probes (Fig. 1B). In each animal, the probe was implanted such that both the pallial structure LMAN and the striatal structure Area X within the avian BG (Person et al., 2008) were targeted (Fig. 1C).

We observed that during putative sleep, slow wave delta activity (1-4 Hz) in the LFP was widespread throughout the pallium and BG, spanning more than 4 mm (Fig. 1D, black arrowheads). Neural spiking was synchronized with the slow wave dynamics and was observable as sparsely bursting patterns of activity in both pallial and BG neurons (Fig. 1D- Spiking Activity). These slow wave dynamics and periods of synchronized bursting were entirely absent from awake data (Fig. 1 E).

We used two complementary approaches to further analyze putative sleep states: 1) we analyzed sleep as a continuous variable using the LFP δ/γ power ratio, and 2) we categorized sleep into discrete stages using an automated avian sleep clustering algorithm (Canavan & Margoliash, 2020; Low et al., 2008). Whereas the former approach acknowledges the continuous nature of sleeping brain dynamics, the latter approach allowed us to numerically characterize certain aspects of avian sleep and compare our results with other avian sleep studies.

To analyze the continuous nature of sleeping brain dynamics, we calculated the δ/γ ratio as the ratio of delta activity (δ, 1-4 Hz) to gamma activity (γ, 30-90 Hz). This method is analogous to metrics that have been used with rodents (Csicsvari et al., 1998; O’Keefe, 1976), but accounts for the dominant frequency bands present during avian sleep (Low et al., 2008; Yeganegi et al., 2019; Yeganegi & Ondracek, 2023).

We calculated the δ/γ ratio for a large range of pallial and BG sites (Fig. 2A). Whereas δ/γ ratios calculated for LFPs within the BG were mostly flat (Fig. 2A, blue lines), δ/γ ratios calculated for pallial LFPs were large and dynamic (Fig. 2A, pink lines, also black line).

Within the pallium, we observed that LPF traces with low δ/γ values had REM sleep-like characteristics (i.e., low amplitude LFP signals), while LFP traces with high δ/γ had SWS-like characteristics (i.e., large amplitude LFP signals; Fig. 2B). Median δ/γ values (Fig. 2C) were calculated using a centrally located pallial LMAN LFP for each bird individually and showed some variability across birds that matched the inter-individual variability reported previously using similar metrics (Yeganegi & Ondracek, 2023). δ/γ values calculated using a LMAN LFP were generally higher than δ/γ values reported using a hyperpallial EEG electrode (Yeganegi & Ondracek, 2023). We used the centrally located LMAN channel quantified in Fig. 2C for all subsequent analysis using the δ/γ ratio.

In addition to the δ/γ ratio, we used a clustering-based sleep scoring method (Canavan & Margoliash, 2020; Low et al., 2008) to classify sleep stages into 3 states (see Methods for more details). For this analysis, we used the same centrally located LMAN LFP that we used in the δ/γ analysis. We observed LFP segments that were classified as SWS were indeed characterized by high δ/γ values, and LFP segments that were classified as REM sleep were characterized by low δ/γ values (Fig. 2D, E) with IS states falling in between the two distributions (Fig. 2F). 3-D plots showed that sleep states were well-segregated on the basis of the spectral features used in the clustering-based approach (Fig. 2G, H). See Fig. S3 for sleep classification distributions for all birds.

### Different spectral features characterize sleep and awake states

Were the activity patterns that we observed during sleep different than those observed during wakefulness? In the next analysis, we examined the spectral features of the LFP signals recorded in LMAN and the surrounding pallium as well as Area X and the surrounding BG during awake and putative sleep phases (Fig. 3A, B). In contrast to the sleep-related activity which was recorded in the dark, the awake recordings took place in the light during the normal day-time hours and captured awake behavior (e.g., singing, moving, resting; Fig. S1).

We observed in LMAN (Fig. 3C, dark gray shading) and the surrounding pallial channels (Fig. 3C, light gray shading) that the power was higher for a broad range of low frequencies during sleep compared to awake states (Fig. 3C, 1-4 Hz, 4-12 Hz, 12-30 Hz, black arrowheads, thin line compared to thick line). In contrast, for LFPs recorded within Area X (Fig. 3C, dark green shading) and the surrounding BG (Fig. 3C, light green shading) only the power in the theta range (Fig. 3C, 4-12 Hz) was occasionally higher during putative sleep compared to awake states (Fig. 3C, black arrowhead, thin line compared to thick line; see also Fig. S4 for data from other birds).

The power in the high-frequency gamma range (Fig. 3C, 90-145 Hz) was higher for LMAN LFPs during awake states compared to sleep states in all birds (Fig. 3C, black arrowhead, see also Fig. S4).

Across depths, we saw that putative sleep states were associated with higher power in low frequency bands compared to awake states (Fig. 3D), which is in line with results from brain areas including RA (Hahnloser et al., 2006) and Area X (S. Y. Yanagihara & Hessler, 2012). Importantly, the LFP associated with the LMAN depth (Fig. 3D, black line; 2.6 mm) was strongly modulated by sleep. We saw that LFPs at the deepest part of the probe followed the common 1/f power spectrum but generally carried less power (Fig. 3D, light green line: 0.1 mm from probe tip).

### Within-area coherence is high during SWS in LMAN and Area X

Given the clear spectral differences that we observed between LMAN and Area X during sleep and awake states, we focused our next analysis on investigating the functional connectivity between these brain areas. We characterized the LFP coherence 1) within LMAN, 2) within Area X, and 3) across LMAN and Area X for 3-s SWS and REM sleep LFP segments and for different canonical frequency ranges that have been shown to play an important role in behavior (Buzsáki & Draguhn, 2004) for all birds (Fig. 4A, B).

**Figure 4:**
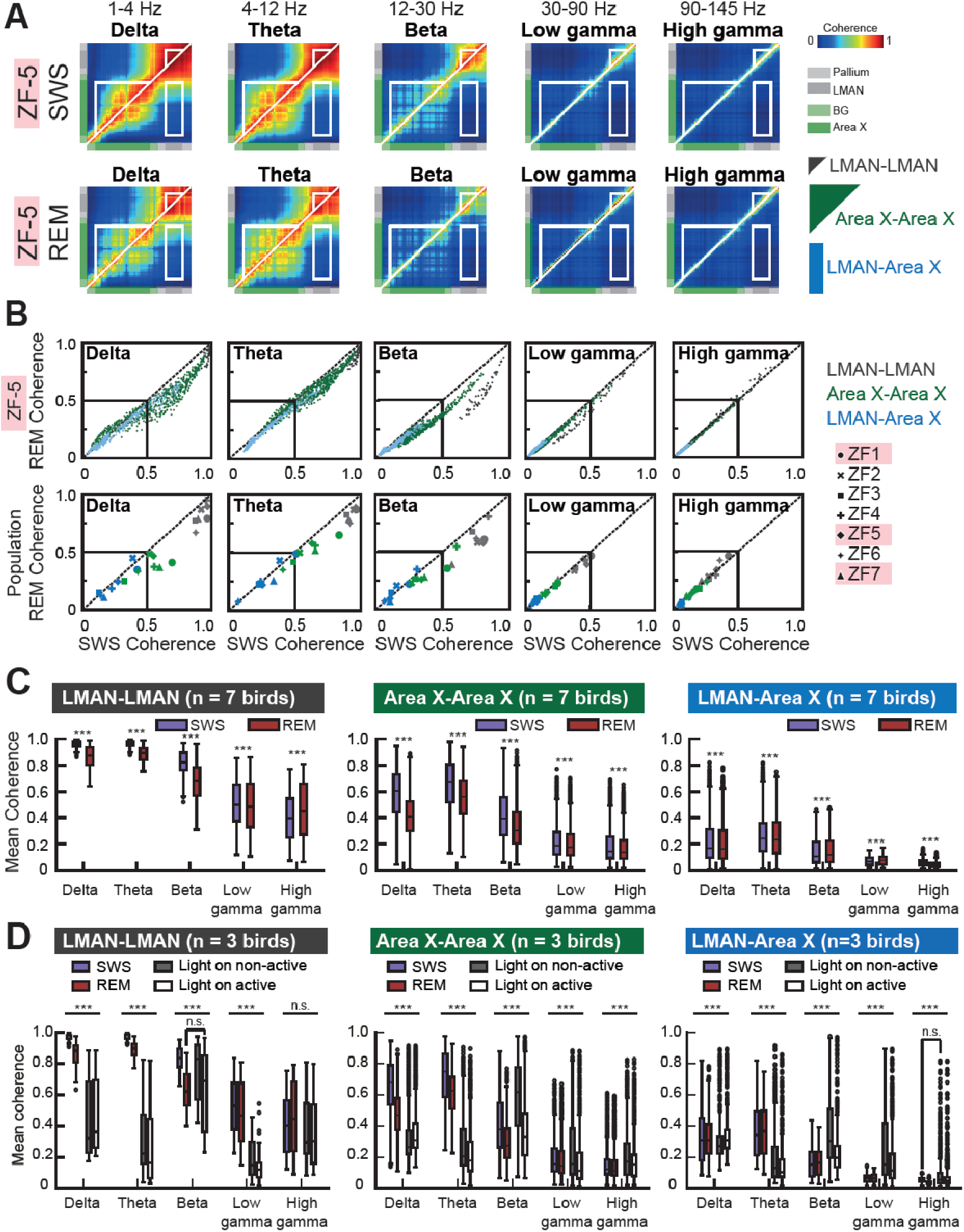
LFP coherence in the pallial-striatal circuit is modulated by behavioral states. (**A)** Heat maps represent a color-coded representation of the average pairwise coherence calculated for all passbands for all SWS segments (top) and REM sleep segments (bottom). Warm colors indicate high coherence values, and cool colors indicate low coherence values. The diagonal white line indicates symmetry. The white triangles along the line of symmetry indicate the pallial electrode sites that were included in the LMAN-LMAN coherency analysis (corresponding to the gray LMAN shading; small black triangle, right side) or the BG electrode sites that were included in the Area X-Area X coherence analysis (green Area X shading; large dark green triangle, right side). The white rectangle (lower right quadrant) indicates the sites that were included in the LMAN-Area X coherency analysis (blue rectangle, right side). Data from sub-adult ZF-5. **(B)** Scatter plots (top row) depict the average REM sleep versus SWS coherence value for each electrode pair plotted for different frequency bands in each panel for sub-adult ZF-5. LMAN-LMAN comparisons (gray dots); Area X-Area X comparisons (green dots) and LMAN-Area X comparisons (blue dots). Black lines at 0.5 for visualization purposes only. Bottom row depicts the mean coherence values for all 7 birds (symbols); colors indicate the coherence comparisons (e.g., LMAN-LMAN in dark gray). Subadult birds are indicated with pink shading of their names. **(C)** Comparison between REM and SWS coherences for different passbands. Box plots indicate the median coherency values across different frequency bands for within-LMAN comparisons (left panel), within-Area X comparisons (middle panel) and LMAN-Area X comparisons (right panels). Blue boxes indicate SWS, red boxes indicate REM sleep. The box bottom and top edges represent the 25th and 75th percentile, respectively, and the middle line represents the median. Whiskers extend to the most extreme datapoints not considered outliers, and datapoints beyond this range are indicated as circles. Asterisks indicate significance (Wilcoxon sign rank test; ***p ≤ 0.001, exact p values are stated in Table 2). **(D)** Comparison between awake and sleep coherences for different passbands for 3 birds. SWS (blue box); REM sleep (red box); Light on non-active (gray box); Light on active (white box). Delta and theta band coherences were significantly lower during the awake states compared to the sleep states for all channel comparisons. Figure conventions same as in (C). Asterisks indicate significance (Friedman test for multiple comparisons; ***p ≤ 0.001, exact p values are stated in Table 3). The non-significant comparisons from the post-hoc Nemeyni test are marked in the figure (n.s). All other pairwise comparisons were significant with the post-hoc test.

**Table 3:**
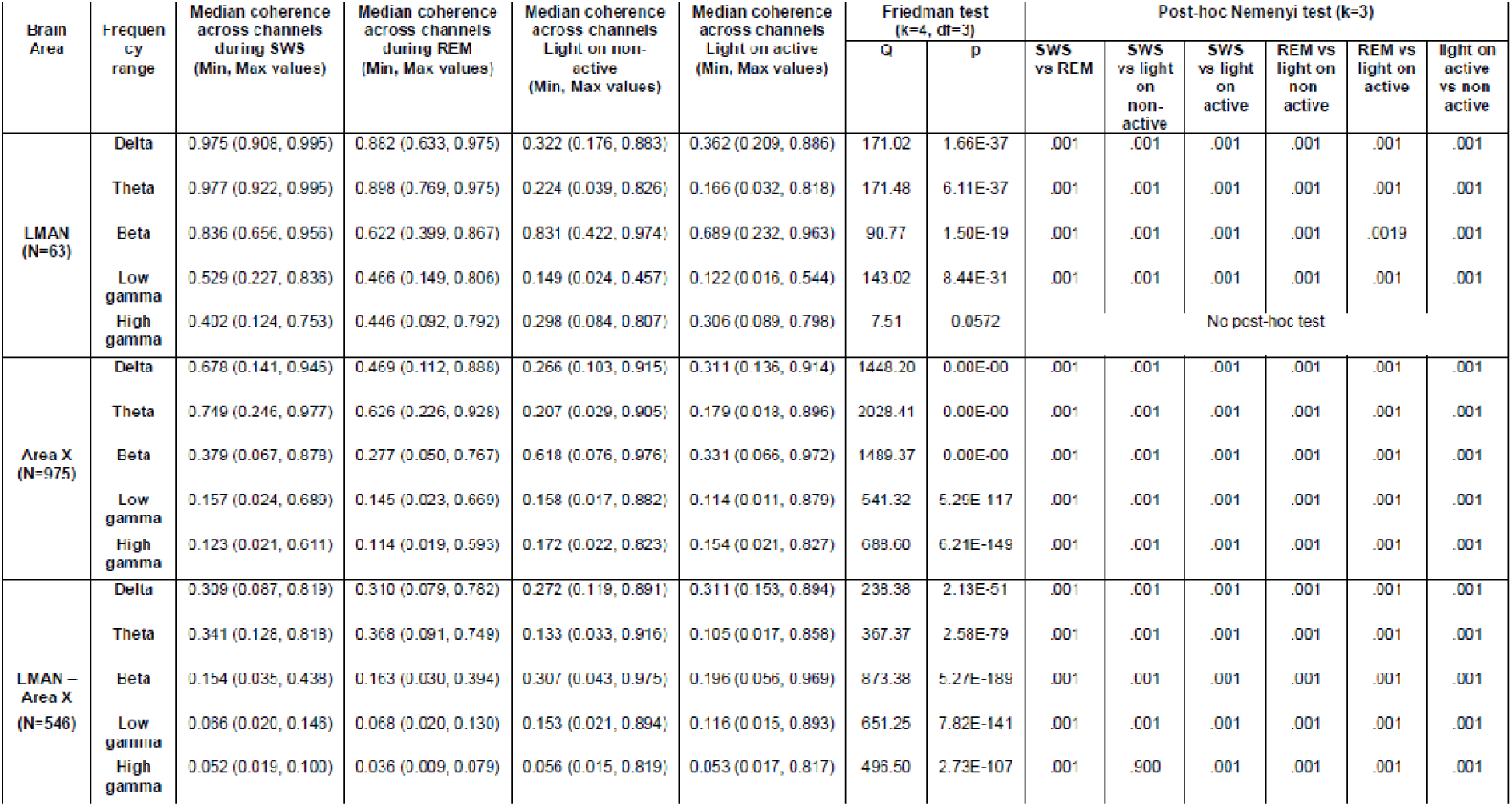
Average coherence values for all birds (n = 3) used for analysis in. **Figure 4D**. Frequency ranges: Delta (1 – 4 Hz), Theta (4 – 12 Hz), Beta (12 – 30 Hz), Low gamma (30 – 90 Hz), High gamma (90 – 140 Hz). N corresponds to the number of LFP pairs.

We observed the average pairwise LFP coherence across birds to be significantly modulated by sleep states (Fig. 4B, Fig. 4C). Whereas coherence in the low frequency ranges was spatially widespread during SWS (Fig. S5), gamma band coherence was limited to spatially neighboring channels with the coherence values dropping to 50% of its peak value within 150 microns. This is consistent with the idea that gamma oscillations reflect local network dynamics (Buzsáki & Wang, 2012; Cardin, 2016; Cardin et al., 2009).

Since the pairwise LFP coherence varied as a function of the distance between the electrode sites and the total number of electrodes within brain regions varied slightly across birds due to variations in implant angle or exact implant co-ordinates, we compared the same number of LMAN and Area X electrodes sites across birds to compare the median coherences between sleep states. The sites were selected such that they spanned the same distance from the identified centers of these brain areas for all 7 birds (see Methods for more details).

We found that SWS and REM sleep coherence values were significantly different for all frequency bands analyzed and across all brain areas (Fig. 4D; Table 2). During SWS within LMAN, we observed very high delta band coherence, ranging from 0.86 - 0.99 (min - max, median = 0.96; Table 2). During REM sleep within LMAN, the delta band coherence was still high (median = 0.87), but the range of values was larger (0.63 - 0.99; min - max), indicating a decrease in coherence between some LMAN pairs during REM sleep.

Within Area X, median coherence values were lower for all frequency ranges and sleep states compared to within LMAN coherence values (Fig. 4B, C; Table 2). Median coherence values within Area X were lower during REM sleep compared to SWS for all frequency ranges (Fig. 4C, Fig. 4B green dots). This finding is consistent with primate literature demonstrating significantly lower coherence within basal ganglia compared to cortical areas during SWS (Mizrahi-Kliger et al., 2018).

LMAN sends direct projections to Area X, which in turn sends feedback projections to LMAN through thalamic area DLM (Fig. 1A; (Farries et al., 2005)). If the closed-loop structure of the circuit plays an important role in modulating sleep dynamics, we may expect strong cross-area coherence.

While median coherence values were low across frequency bands for LMAN-Area X LFP pairs compared to within-LMAN or within-Area X LFP pairs (Table 2), a few individual LMAN-Area X LFP pairs showed high coherence values > 0.5, especially for the low frequencies (1-12 Hz; Fig. 4B, blue dots). This finding suggests that the input from LMAN to Area X may be spatially restricted, as only a few pairs shared similar oscillatory drive during sleep.

We did not find a strong effect of sleep state on the across-area LMAN-Area X coherence. Although significant differences were found across all frequency bands (Fig. 4C, right), the median differences between SWS and REM sleep were quite small (Table 2).

### LMAN-Area X dynamics are modulated by behavioral state

We were curious as to how the LFP coherence changed as an effect of behavioral state, and so we compared the sleep-related LFP coherence with coherence analyzed for daytime “Light on active” and “Light on non-active” periods for a small group of birds (n=3 birds; Fig. 4D). The sleep-related coherence analysis for this smaller group of birds was consistent with the results of the larger group of seven birds (see Table S2): coherence values were highest within LMAN compared to within Area X and across LMAN and Area X pairs for all frequency bands, and coherence values were higher during SWS compared to REM sleep across frequency bands, with the exception of high gamma within LMAN. In this case, REM coherence was higher compared to SWS coherence within LMAN; however, this difference was not significant for the smaller group of three birds (Fig. 4D, LMAN-LMAN comparison; High gamma, n.s.).

When we compared sleep-related coherence to daytime coherence (i.e., Light on active and Light on non-active), we saw that the delta and theta band coherence values were significantly lower for daytime periods, suggesting that periods of putative sleep in the dark (i.e., SWS and REM states) were distinct from periods of active and non-active behaviors recorded during the daytime.

Interestingly, when we analyzed the coherence trends within the daytime periods, we observed that periods of daytime active behavior were associated with higher delta band coherence values compared to non-active periods of daytime. This was true for within LMAN, within Area X, and across LMAN-Area X LFP pairs, suggesting that oscillatory activity in the AFP differs during non*lz*active daytime periods compared to nighttime sleep

### Phases of high δ/γ activity during sleep are associated with increased spiking variability for neurons in LMAN but not Area X

Using our Neuropixels recordings, we leveraged the high-pass filtered electrode signals to investigate the spiking activity in LMAN and Area X during sleep in n = 5 birds (Table 1). Spikes were identified and assigned using state-of-the-art automatic detection and sorting algorithms, either in a fully automated way or followed by meticulous manual curation to ensure accuracy and reliability (see Methods). The results of any spiking analysis did not differ qualitatively across these two preprocessing approaches.

During sleep, we observed increased but sparse population-wide spiking activity which was locked to negative LFP deflections and interleaved with periods of neural silence, followed by sudden switches to periods of random spiking and lower LFP amplitudes (see Fig. 1D). To quantify the different patterns of spiking activity that we observed during sleep, we calculated the coefficient of variation (CV, standard deviation divided by the mean) for identified LMAN and Area X neuron populations in 10 s bins.

The CV can be thought of as a measure of collective variability in coincident spike trains: CV values for the population rate will approach zero when all neurons are statistically independent (Kobak et al., 2019). This is because random irregularities stemming from individual neurons will average out as the population size increases, ultimately leading to a stable population rate. However, when neurons are synchronized and activity oscillates, the variability from one moment to another in the population rate is high.

We found that median CVs ranged from 0.27 in Area X to 2.14 in LMAN (Fig. 5A; see Table 4 for individual values), and CV values were significantly higher for LMAN neurons compared to Area X neurons in all birds (median [min, max] LMAN CV over all birds = 1.44 [0.73, 2.14]); median [min, max] Area X CV for all birds was = 0.61 [0.27, 1.04]).

**Figure 5:**
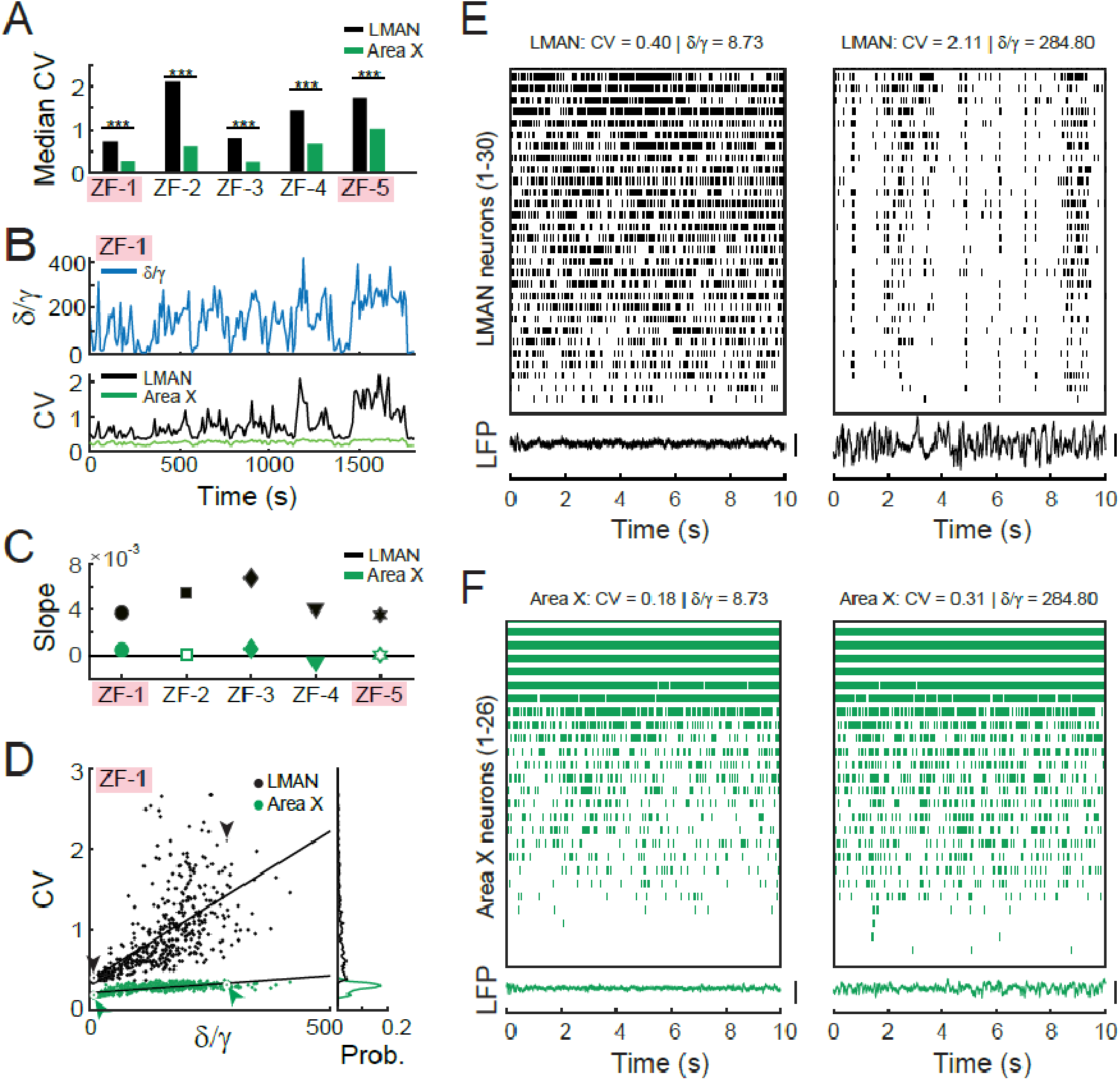
Spiking variability is significantly higher for LMAN neurons compared to Area X neurons. **(A)** Bar plots summarize CV values for 7 finches (black bars, LMAN population; green bar, Area X population). CV values were higher for LMAN populations compared to Area X populations. Sub-adult birds are indicated by pink shading of their names. **(B)** δ/γ values (top, blue line) were highly correlated with CV values (bottom) for both LMAN and Area X populations. δ/γ values and CV values computed in 10 s windows. **(C)** The slope of regression fits between CV and δ/γ was consistently higher in LMAN compared to Area X across birds. Filled symbols indicate significant correlations (p < 0.01, Table 5). **(D)** Scatter plot indicates the CV versus δ/γ values for ZF-1. LMAN, black dots; Area X, green dots. Black and green arrows indicate the data points that correspond to the 10 s snippet of data plotted in (D) and (E). The marginal distributions for the CV values are plotted (right) for LMAN (black line) and Area X neurons (green line). **(E)** Spiking raster plot for 33 LMAN neurons for an example time window with low CV value (left plot) and a high CV value (right box). Each tick corresponds to a spike and neurons are sorted by their average spiking activity from high (top) to low (bottom). Corresponding 10 s LFP trace is plotted at the bottom of the raster plot. Vertical bar is 500 uV. Data correspond to black arrows in (D). **(F)** Spiking raster plot for 27 Area X neurons for a low CV value (left plot) and a high CV value (right box). Figure convention same as in (E). Data correspond to green arrows in (D). Vertical bar is 500 uV.

**Table 4:** Median CV values were significantly higher for LMAN neurons compared to Area X neurons. Median values and range. Population statistics are computed across median CVs. p value is the result of a Wilcoxon rank sum test.

| Bird | N | LMAN CV median (min, max) | Area X CV median (min, max) | Wilcoxon ranksum test |  |
| --- | --- | --- | --- | --- | --- |
|  |  |  |  | z Value | p Value |
| ZF-1 | 532 | 0.73 (0.33-2.68) | 0.28 (0.14-0.37) | 28.18 | 1.08E-174 |
| ZF-2 | 315 | 2.14 (0.32-5.16) | 0.62 (0.39-0.99) | 18.13 | 1.74E-73 |
| ZF-3 | 990 | 0.80 (0.39 -6.42) | 0.27 (0.16-0.54) | 38.12 | 6.71E-318 |
| ZF-4 | 318 | 1.44 (0.44-5.44) | 0.67 (0.43-0.98) | 15.79 | 3.94E-56 |
| ZF-5 | 400 | 1.62 (0.88-5.08) | 1.04 (0.86-1.22) | 18.36 | 3.00E-75 |
| Population median | 5 | 1.44 | 0.62 |  |  |

**Table 5:** LMAN CV values were significantly and positively correlated with δ/γ values for 5/5 zebra finches. Correlations were more heterogeneous for Area X. R, Pearson correlation coefficient.

| Bird | | LMAN: CV versus $\delta/\gamma$ | | | Area X: CV versus $\delta/\gamma$ | |
| --- | --- | --- | --- | --- | --- | --- |
|  | N | R | p |  | R | p |
| ZF-1 | 532 | 0.67 | 7.58E-71 |  | 0.73 | 2.45E-88 |
| ZF-2 | 315 | 0.43 | 2.08E-15 |  | 0.05 | 0.42 |
| ZF-3 | 990 | 0.32 | 1.17E-25 |  | 0.24 | 2.06E-14 |
| ZF-4 | 318 | 0.16 | 0.0034 |  | -0.28 | 3.65E-07 |
| ZF-5 | 400 | 0.71 | 5.40E-63 |  | -0.09 | 0.085 |

We observed that the CV values fluctuated closely with δ/γ values, such that high CV values were associated with high δ/γ values (Fig. 5B). This trend was especially obvious for LMAN neurons (Fig. 5C, D). Indeed, regression fits between CV values for LMAN neurons and δ/γ values returned higher slopes (Fig. 5C, D) and both variables were highly correlated for all birds (Table 5). This was not the case for Area X neurons: estimated regression slopes were lower, and only 2/5 birds showed significant positive correlations between CV values and δ/γ values (ZF-1 and ZF3), whereas 1 bird showed a significant negative correlation (ZF-4, Table 5; Fig. 5C, filled symbols).

High CV values for LMAN neurons compared to lower CV values for Area X neurons (Fig. 5A) suggest that the spike trains for LMAN neurons were more irregular than for Area X neurons. Indeed, when we examined spike rasters corresponding to high CV values for LMAN populations (Fig. 5D, E), spiking patterns were conspicuously irregular - albeit largely synchronized as bursts of spikes across the population of LMAN neurons. In contrast, CV values were significantly lower for Area X neurons (Fig. 5A), capturing their regular spiking patterns and high firing rates (Fig. 5D, F).

It is important to note that the δ/γ trace used in this analysis was calculated from a centrally located LMAN LFP (see Methods for details). Therefore, strong correlations between CV values and δ/γ values in LMAN may reflect the tight correspondence between spiking activity and LFP oscillations in LMAN. The correspondence between spiking activity in Area X and the LMAN LFP was much more heterogeneous, suggesting that oscillatory activity in LMAN did not directly influence Area X spiking activity during sleep as it did in LMAN.

### Synchronized population bursting in LMAN during sleep is associated with a negative deflection in the LFP

Our findings suggested that within LMAN, populations of neurons are co-activated at specific time points during putative sleep. In order to explore the synchronized bursting of LMAN neurons during putative sleep - and how they are related to the LFP (Fig. 6A) - we identified and thresholded bursting events in the population of LMAN neurons (Fig. 6B). Then, we used these synchronized bursts to identify activity patterns in the LFP, which we called “burst-aligned LFP” (Fig. 6C). Individual burst-aligned LFP segments generally had higher power in frequency ranges < 10 Hz (example in Fig. 6D), and the average burst-aligned LFP showed a prominent low-frequency power burst centered on zero-time lag (Fig. 6E). Indeed, when we compared the bursting-aligned LFP segments (Fig. 6C, lower blue line) to randomly selected LFP segments (Fig. 6C, lower black line), the bursting-aligned LFP segments showed a notable increase in power for frequencies lower than 10 Hz. This phenomenon was present across birds: every bird examined had a notable negative deflection that corresponded to the population synchronized bursting (Fig. 6F).

**Figure 6:**
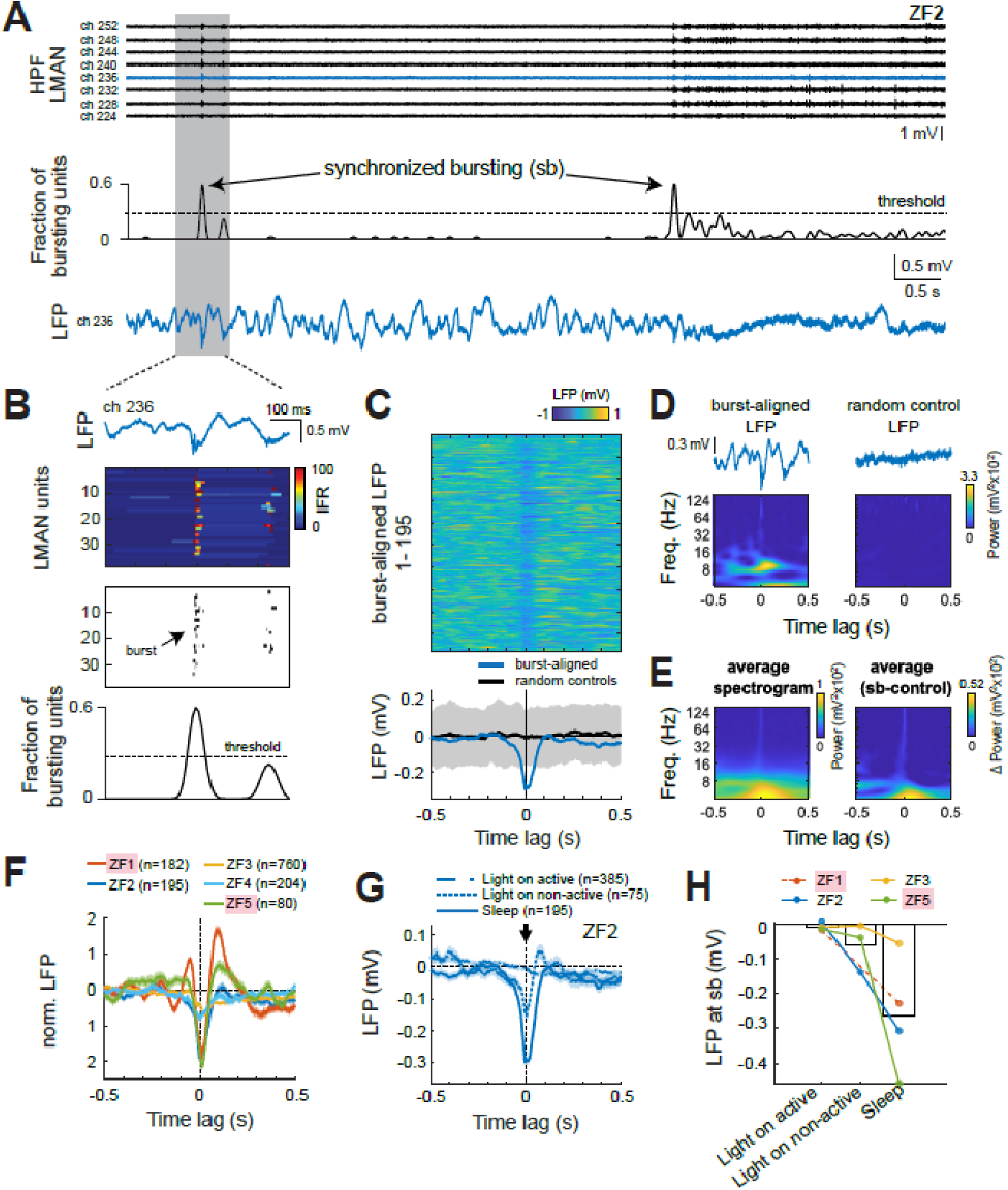
Synchronized bursting in LMAN is associated with a negative deflection in the LFP. **(A)** Two examples of synchronized bursting activity in LMAN from ZF-2. Top panel depicts high-pass filtered (HPF) electrode signals from eight electrodes in LMAN. Middle panel depicts the detection of synchronized bursting (black line) based on the smoothed fraction of bursting units in a 50 ms window exceeding the 99.5^th^ percentile threshold. Bottom panel depicts the raw LFP (blue, same electrode as in top panel). Area indicated by gray shading is expanded in (B). **(B)** Zoom-in on the highlighted gray window in (A). Top panel depicts the instantaneous firing rate (IFR) of all LMAN units (n = 32), color-coded and clipped at 100 Hz. The panel below depicts the occurrence of bursts in black for each LMAN unit. Lower panels same as in (A). **(C)** Upper panel depicts 195 bursting-aligned LFP segments detected from channel 236, stacked and color-coded. Events are sorted by the total fraction of bursting units. Lower panel indicates the average bursting-aligned LFP (blue) and the same statistics for randomly selected LFP segments (black, gray shading is +/- SD). **(D)** Two raw LFP snippets (top panel) and their corresponding scalograms (bottom panel). The left example shows a bursting aligned LFP. The right depicts random control (right). **(E)** The average bursting aligned-LFP scalogram (left) and its difference to the average random control scalogram (right). Note how the power density for the synchronized burst is highly elevated at low frequencies. Sb, synchronized burst. **(F)** LFPs show consistent negative dip during a synchronized burst. Each line indicates the bursting-aligned LFP normalized to the random control (z-norm ± SEM) for each bird. **(G)** The average burst-aligned LFP ± SEM across different states for ZF-2. Note that the LFP dip remains visible during light on non-active periods (short-dashed line), though it is reduced in magnitude, and becomes further diminished during light on active periods (long-dashed line). **(H)** Comparison of LFP dips across all birds with available awake data shows that the strongest deflections occur during light-off periods. No bursting events were detected for ZF-1 during light on active periods. See Fig. S6 for more details.

Having established that synchronized bursting in LMAN during putative sleep is consistently associated with prominent low-frequency deflections in the LFP, we next asked whether similar events occur during wakefulness. Specifically, in the four birds for which we recorded neural activity during daytime periods, we examined whether synchronized LMAN bursting events also occur during active and resting periods, and if so, whether the associated burst-aligned LFP exhibits features similar to those observed during putative sleep.

During light on non-active periods, burst-aligned LFPs in ZF-2 and ZF-3 resembled those seen during putative sleep, exhibiting a negative deflection in the LFP at the time of the synchronized bursting and elevated low-frequency power in the scalogram, although with reduced overall magnitude (Fig. 6G; Fig. S6). In contrast, ZF-1 exhibited no detected events during rest, and ZF-5 showed only 11 events, none of which were associated with a clear LFP deflection.

In active light on periods, we observed numerous synchronized bursting events across all birds except ZF-3, which was generally inactive during this recording (Fig. S1). The detected events coincided with behaviors such as singing, calling, or general movement, consistent with the increased bursting activity characteristic during vocal production (Hessler & Doupe, 1999b). However, the associated LFPs remained flat across all birds, showing no consistent deflection time-locked to the bursts (Fig. 6G, long dashed line; Fig. S6). This absence of coordinated low-frequency structure in the LFP distinguishes these active-state bursting events from those observed during putative sleep, where synchronized bursts are reliably coupled with prominent negative LFP deflections (Fig. 6F).

Consistent with these observations, Fig. 6H summarizes the average LFP amplitude at the time of synchronized bursts across states, showing the strongest negative deflections during putative sleep, weaker deflections during light on rest, and values near zero during active wakefulness. This graded pattern supports a state-dependent modulation of LFP responses to synchronized bursting in LMAN.

### Synchronized population bursts in LMAN occur predominantly during NREM sleep

We were curious as to whether these synchronized bursting events in LMAN occurred during a specific sleep regime, i.e. either during NREM sleep (i.e., SWS and IS sleep) or during REM sleep (Fig. 7A). With the exception of 1 bird (Fig 7A, ZF-4: upward triangle), the synchronized bursting events occurred predominantly during NREM sleep.

**Figure 7:**
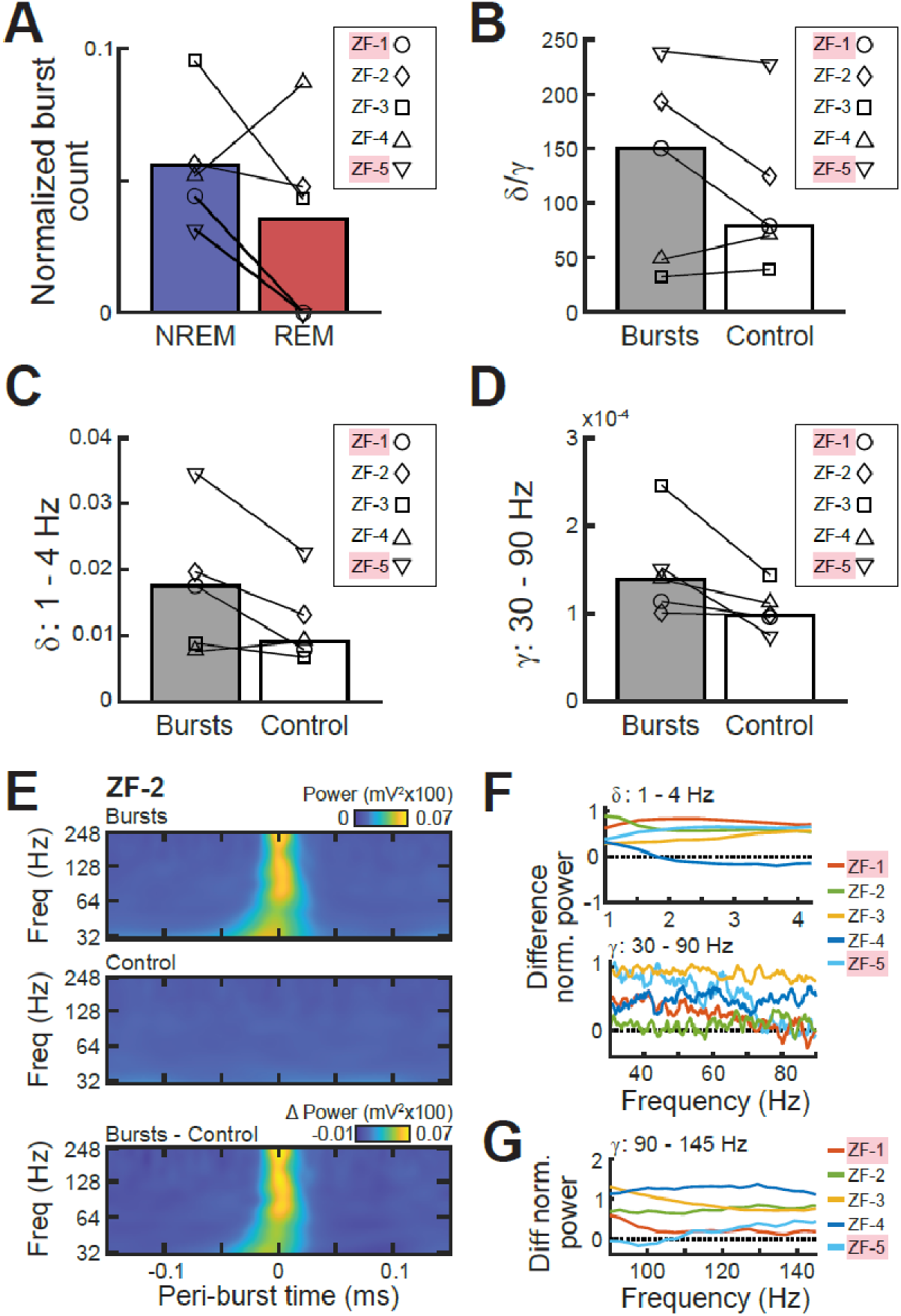
Synchronized bursts occur during NREM sleep and are associated with higher γ activity. **(A)** Bar plots indicate median number of bursts associated with each sleep state normalized by the overall sleep state duration in seconds across birds (see Fig. 2F, G). With the exception of ZF-4, most birds showed more bursts detected during NREM states compared to REM sleep states. NREM states included both SWS and IS classified states. Symbols indicate median data for each bird. Sub-adults are indicated with pink shading of their names. **(B)** Bar plots indicate median δ/γ value calculated for 500 ms segments of LFP that were centered on a synchronized burst detection (Bursts, grey bar) compared to the mean value of 500 ms control segments of LFP that were temporally shifted away from the burst detection by a random offset (Control, white bar; see Methods for details). Symbols indicate median data for each bird. Note how δ/γ values are on average higher for the control LFP segments. **(C)** Same convention as in (B) for δ values. Note how δ values are higher for the burst LFP segments compared to the control LFP segments. **(D)** Same convention as in (B) and (C) for γ values. Note the higher γ values during the burst LFP segments compared to the control LFP segments. **(E)** Spectrograms display the average peri-burst time window surrounding a burst event (top), a control event (middle), and the difference in power between the burst event and the control event (bottom) for ZF-2. Note the increase in power surrounding the burst event that extends up into the high γ range. **(F)** Panels display the normalized PSD for different frequency ranges: δ (1-4 Hz, top), low γ (30-90 Hz, middle), and high γ (90-145 Hz, bottom plot). Normalized PSD indicates the difference between the burst events and the control events divided by the standard deviation of the control events. Different line colors indicate different birds. Note how data for all birds fall above the zero (dotted line) indicating higher values during the burst event compared to control events.

We also examined the δ/γ values surrounding the synchronized bursting events, with the expectation that these events might occur during LFP segments with higher δ/γ values (i.e., during SWS). Interestingly, we found that the bursting events were not consistently associated with lower δ/γ values compared to control events across birds (Fig. 7B; see Methods for details). Because the δ/γ is a ratio, low δ/γ values can result from a decrease in δ power or an increase in γ power. We examined the δ and γ power separately for bursting events compared to control events. We found that while the power in the δ range was indeed elevated during bursting events compared to control events for all but one bird, ZF-4, (Fig. 7C), a larger increase in γ power during the bursting event (Fig. 7D) compared to control segments led to the overall inconsistent δ/γ values we observed during bursting events compared to controls (Fig. 7B).

When visualized as the difference between the burst and control segments (Fig. 7E), this burst-associated increase in power extended from delta up through high gamma and was present for all birds analyzed (Fig. 7F).

## Discussion

In order to explore the ongoing spontaneous activity patterns of the AFP during sleep and awake behavior in songbirds, we investigated the activity patterns of neurons in LMAN and Area X (Fig. 1A, C) using chronically implanted Neuropixels probes (Fig. 1B).

Coherence between field potentials has been used in a multitude of studies as a mesoscopic measure of functional connectivity between brain areas (Schneider et al., 2021) and is believed to reflect the neuronal connectivity (Buzsáki & Schomburg, 2015) and oscillatory synchronization (Bastos et al., 2015) in these brain areas.

Importantly, we observed strong significant differences between sleep-related coherence (i.e., SWS and REM) and awake-related coherence (i.e., light on non-active and light on active) across brain regions and passbands (Fig. 4D, see exact values in Table 3). This finding demonstrates that the birds were in a different behavioral state in the dark compared to recordings in the light, and suggests that our sleep recordings were not overly contaminated with undetected quiet awakenings. If this had been the case, we hypothesize that the sleep-related coherence values would more strongly resemble the light on non-active coherence values, which they do not (see Text S1 for further discussion).

Contrary to our expectations, we did not observe strong across-area coherence between LMAN and Area X on average, although individual LMAN-Area X pairs did show strong coherence > 0.5. In contrast, studies of motor skill learning in rodents found an increase in cross-area coherence between motor cortex and striatum during sleep (Koralek et al., 2013; Lemke et al., 2021). The fact that we did not observe strong cross-area coherence may stem from the fact that the birds in this study were not actively engaged in a learning paradigm.

A previous study demonstrated the presence of phasic LFP gamma oscillations in Area X during sleep, with singing-modulated Area X units phase-locked to gamma oscillations (S. Y. Yanagihara & Hessler, 2012). Our pairwise LFP coherence results, albeit a mesoscale measure of synchronization across brain areas, could be a reflection of this phase-locking and the concerted activity of anatomically connected LMAN units and Area X units during awake periods. Verification of this hypothesis would require further data collection in awake and singing birds to investigate the specific neural activity of different neuronal subtypes and their relation to the ongoing LFP gamma oscillations.

During putative sleep, we observed that populations of LMAN neurons spontaneously reactivated in bursting events. These events occurred predominantly during NREM sleep and were associated with a transient increase in gamma band power. Across animals, the population bursting events were preceded by a large negative deflection of the LFP, pointing to the possibility that they were triggered by the ongoing large-scale global oscillatory activity in LMAN.

The combination of large negative LFP deflection and population bursting activity in LMAN is highly reminiscent of SWR activity observed during SWS and quiet rest in rodent hippocampus (Buzsáki, 1986, 2015; O’Keefe, 1976).

The sharp wave and ripple LFP patterns that characterize mammalian SWRs are thought to arise from the specific anatomy of the hippocampus. The sharp wave reflects massive excitation of CA1 neurons by the pyramidal cells in CA3. This excitation synchronizes the interneuron network and generates a ripple in the pyramidal layer (Buzsáki et al., 1983; Girardeau & Zugaro, 2011; A. A. Liu et al., 2022). In rodents, the ripple oscillation is characterized by a frequency in the range of 110-200 Hz, overlapping with typical gamma band activity (Buzsáki, 2015; A. A. Liu et al., 2022).

Our observation of SWR-like activity in LMAN suggests that sharp waves coinciding with gamma oscillations may not be unique to mammalian hippocampus. However, the function of this non-hippocampal SWR-like activity remains unclear.

Hippocampal SWR activity has been typically considered a “biomarker” that accompanies spatial navigation learning and episodic memory (Buzsáki, 2015); these learning paradigms are fundamentally different from the auditory and motor learning that characterizes vocal learning in songbirds. Furthermore, the birds used in our study sang mature crystallized songs characterized by stable motif structure, suggesting that the developmental learning period had already ended. Nevertheless, LMAN has been shown to have an important role in the ongoing maintenance of adult song via N-methyl-D-aspartate (NMDA) receptor-dependent transmission from LMAN to the pallial premotor nucleus RA (Brainard & Doupe, 2000; Solis et al., 2000; Warren et al., 2011). Bursting in LMAN influences RA and thereby affects the spectral variability of song (Kao et al., 2005; Moorman et al., 2021; Tian et al., 2023; Woolley & Kao, 2015). From these observations a potential role of avian SWR activity emerges in the consolidation of experience in a broader sense (Tachibana et al., 2022).

It remains to be determined whether the LMAN synchronized bursts that we observed during sleep represent true replay events related to, for example, daytime singing. “Replay”-like reactivations have also been reported during sleep in RA (Dave & Margoliash, 2000; Elmaleh et al., 2021, 2023). Given the widespread LFP dynamics that we observed across LMAN, it would be important to investigate how synchronized bursting in LMAN and RA are temporally related, and specifically whether the widespread LFP oscillations in LMAN influence RA bursting during sleep.

Our investigation of the spontaneous network activity of the avian AFP has revealed two major insights that help inform our understanding of how the AFP compares to the mammalian corticostriatal circuit.

Similar to what has been reported in primates, we observed an overall desynchronization of activity in BG compared to pallial areas during SWS sleep in songbirds. Specifically, pairwise LFP coherence values within Area X were lower compared to LMAN for low frequency ranges during SWS (Fig. S5: delta, theta, beta). This finding is consistent with primate literature demonstrating significantly lower coherence within BG compared to cortical areas during SWS (Mizrahi-Kliger et al., 2018).

In contrast, the presence of SWR-like activity in LMAN is unexpected and suggests that multiple pallial neural architectures may give rise to SWR-like activity patterns.

During embryonic development in mammals, parts of the ventral pallium give rise to the amygdalar complex (Garcia-Calero & Puelles, 2021). In birds and reptiles, however, the ventral pallium gives rise to the sauropsid brain area known as the dorsal ventricular ridge (DVR; (Puelles et al., 2000, 2017)). This large pallial region dorsal to the BG is unique to reptiles and birds. In birds, the DVR has functionally been compared to the mammalian neocortex due to its role in avian cognition (Güntürkün, 2005): high-level cognition in corvids (Nieder et al., 2020), vocal learning in songbirds (Roberts et al., 2017; S. Yanagihara & Yazaki-Sugiyama, 2016), and visual imprinting in chicken (Horn, 2004) are all behaviors that critically involve the DVR. In birds, the DVR is subdivided into the mesopallium, nidopallium, entopallium, and arcopallium (Briscoe & Ragsdale, 2018); LMAN is part of the nidopallium.

Importantly, similar SWR-like LFP patterns have also been recorded during sleep in the reptilian DVR (Albeck et al., 2022; Libourel et al., 2018; Shein-Idelson et al., 2016). The fact that SWR-like activity patterns arise from both reptilian and avian DVR during sleep suggests that some version of the neural circuit that gives rise to mammalian hippocampal SWRs may also be present in the avian DVR – despite the absence of clear anatomical lamination in the DVR (although see (Stacho et al., 2020)). As highlighted above, however, the function of this DVR-specific, SWR-like activity is still a mystery. Further work is required to determine whether the SWR-like activity observed in the avian DVR is an “epiphenomenon” related to the ongoing dynamics of the sleeping brain, or whether it has a true function in learning and memory.

## Acknowledgements

This research was supported by the Agence Nationale de la Recherche (SleepinBrainDyn) and the CNRS (International Emerging Action, project BirdSleep) to AD; the EUR NEURO (EUR/7410-5) and Bordeaux Neurocampus - GPR BRAIN 2030 (GP/7200R-12) to EGZC; the Deutsche Forschungsgemeinschaft (ON 151/1-1) and the Daimler und Benz Stiftung (PN 32-09/18) to JMO; the BayFrance program (FK23 2019) to CL, HY, NG, and JMO. The authors would further like to thank and acknowledge Melanie Dumont for excellent animal caretaking and Manon Rolland for help with experiments.

