## Supplementary Materials for "Sharp waves, bursts, and coherence: Activity in a songbird vocal circuit is influenced by behavioral state"

Abbreviated title: Pallial-striatal dynamics and behavioral state

Author Names and affiliations: Corinna Lorenz[1,2]*, Anindita Das[2]*, Eduarda Gervini Zampieri Centeno[3,5], Hamed Yeganegi[4,6], Robin Duvoisin[1], Roman Ursu[3], Aude Retailleau[3], Nicolas Giret[2], Arthur Leblois[3], Richard H. R. Hahnloser[1], and Janie M. Ondracek[4†]

[1] Institute of Neuroinformatics, University of Zurich and ETH Zurich, 8057, Zurich, Switzerland
[2] Institut des Neurosciences Paris-Saclay, CNRS, 151 route de la rotonde, 91400 Saclay, France
[3] Bordeaux Neurocampus, Institut des Maladies Neurodégénératives, 146 rue Léo-Saignat - 33076 Bordeaux Cedex
[4] Technical University of Munich, TUM School of Life Sciences, Chair of Zoology, Liesel-Beckmann-Straße 4, 85354 Freising, Germany
[5] Amsterdam UMC location Vrije Universiteit Amsterdam, Anatomy and Neurosciences, De Boelelaan 1117, Amsterdam, The Netherlands
[6] Graduate School of Systemic Neurosciences, Ludwig-Maximilians-University Munich, Großhaderner Str. 2, 82152, Planegg, Germany
(*) Authors contributed equally
(†) Corresponding author

### Supplementary Material

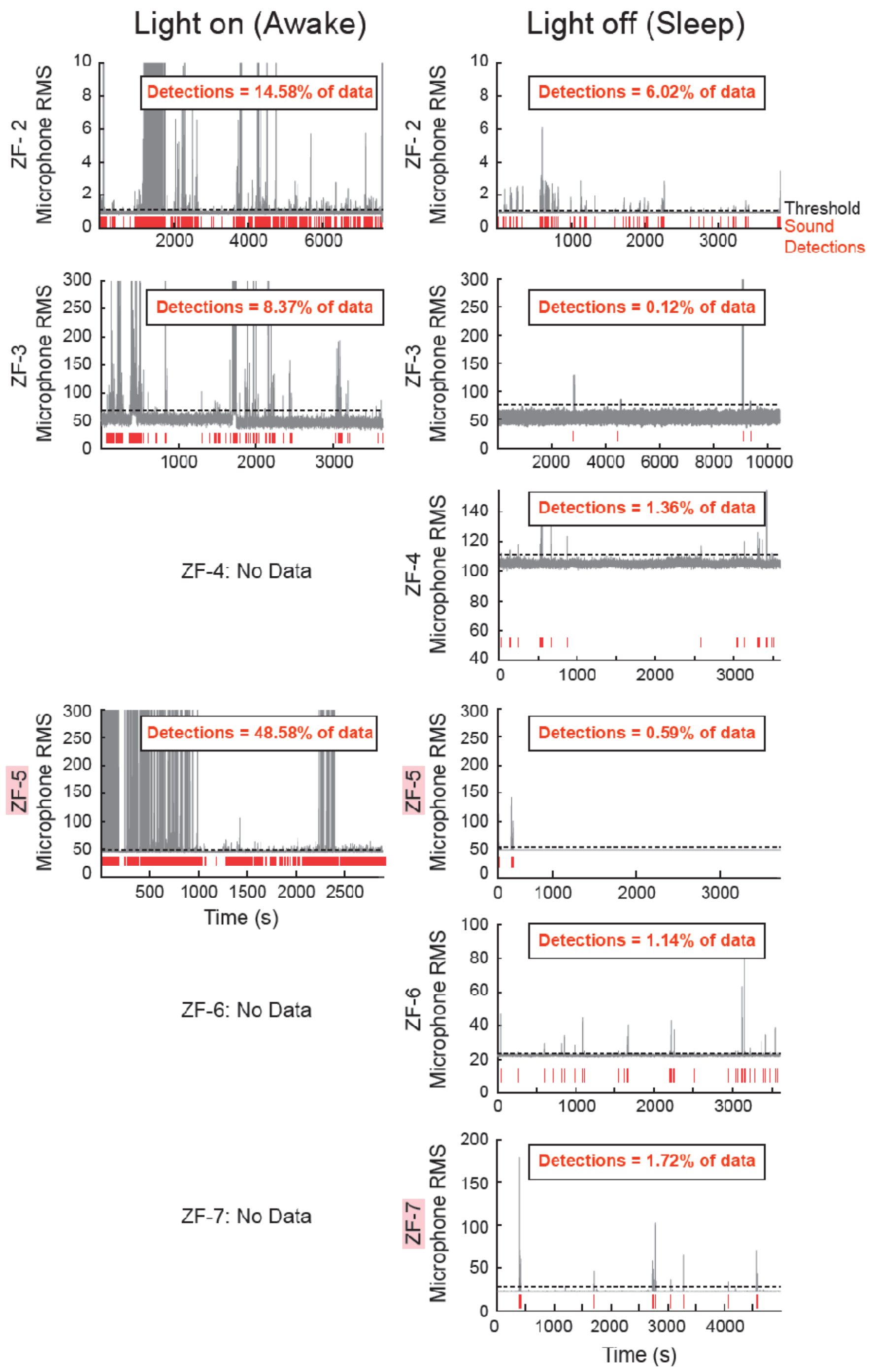


**Supplementary Figure S1: Comparison of light on and light off recordings and corresponding sound detections for all birds** Comparison of microphone signals recorded during the light on (awake) period (left column) and the light off (sleep) period (right column). Gray traces indicate the root mean square (RMS) of the microphone signal. Black dotted line indicates the manually set threshold, and threshold crossings indicate sound detections (red lines). The overall percent of data for which sounds were detected is stated (top). Sub-adult birds are indicated with pink shading of their names. No awake data were record for ZF 4, 6, and 7.

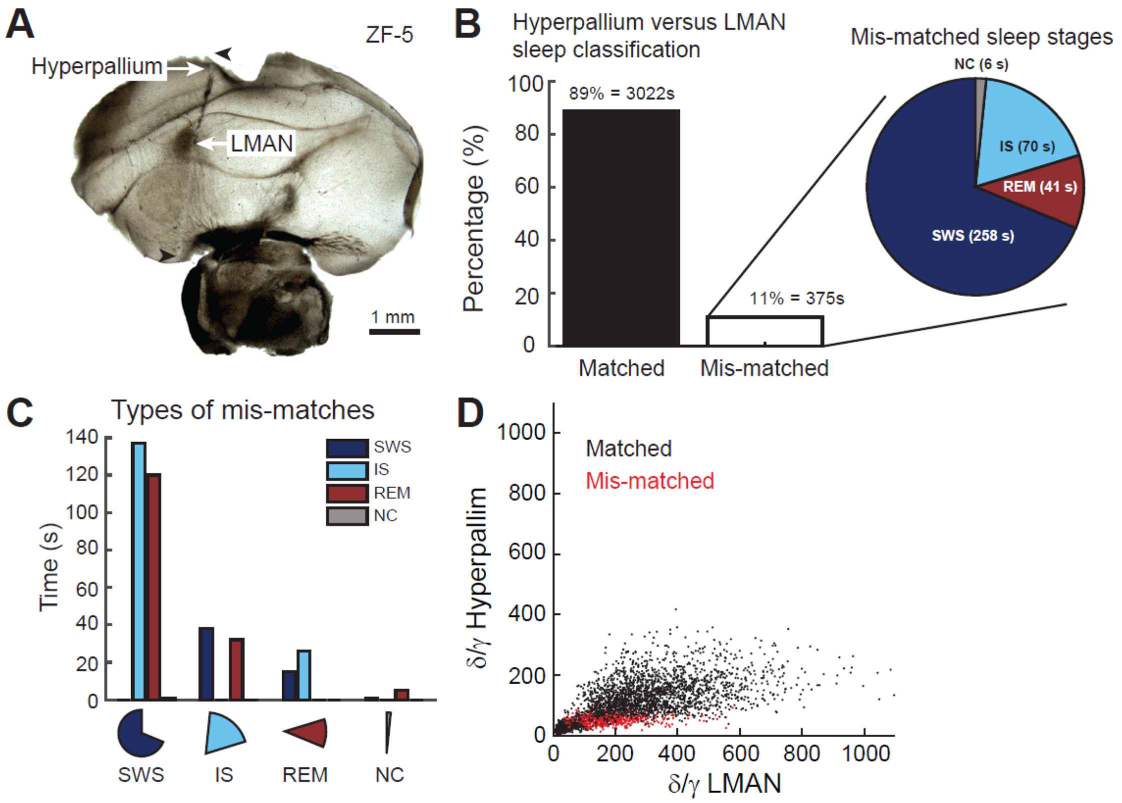


**Supplementary Figure S2: Comparison of sleep stage classification for hyperpallium and LMAN LFPs (A)** Histological brain slice depicts the electrode tract (black arrow heads). The electrode was positioned such that recording sites were available in the upper hyperpallium as well as the deeper LMAN. Histological slice was cut at room temperature on a vibratome from sub-adult ZF-5. **(B)** Bar plot indicates the number of matching sleep stage classifications between the hyperpallium LFP and the LMAN LFP (89%, black bar) and the number of mismatches (11%, white bar). Of the 11% mis-matched sleep stages, the pie chart on the right indicates which LMAN sleep stages did not match the classification of the hyperpallium LFP, e.g., 258 s of SWS in LMAN were classified as a different sleep state in the hyperpallium. The color code represents the sleep stage classification. **(C)** The exact breakdown of the LMAN mismatches is depicted in bar plots. Each bar is color-coded to represent a different sleep state. For the mismatched LMAN SWS states (far left bars), we see that almost 140 s were cases where SWS in LMAN was labeled as IS sleep in the hyperpallium. **(D)** For each 1 s bin of the entire recording, the data were plotted as the d/y ratio for the hyperpallium channel (y axis) versus the d/y ratio for the LMAN channel (x-axis). Matched points are plotted in black, and the mismatched sleep stages are plotted in red. Note how the range of d/y values is much smaller for the hyperpallium channel compared to the LMAN channel.

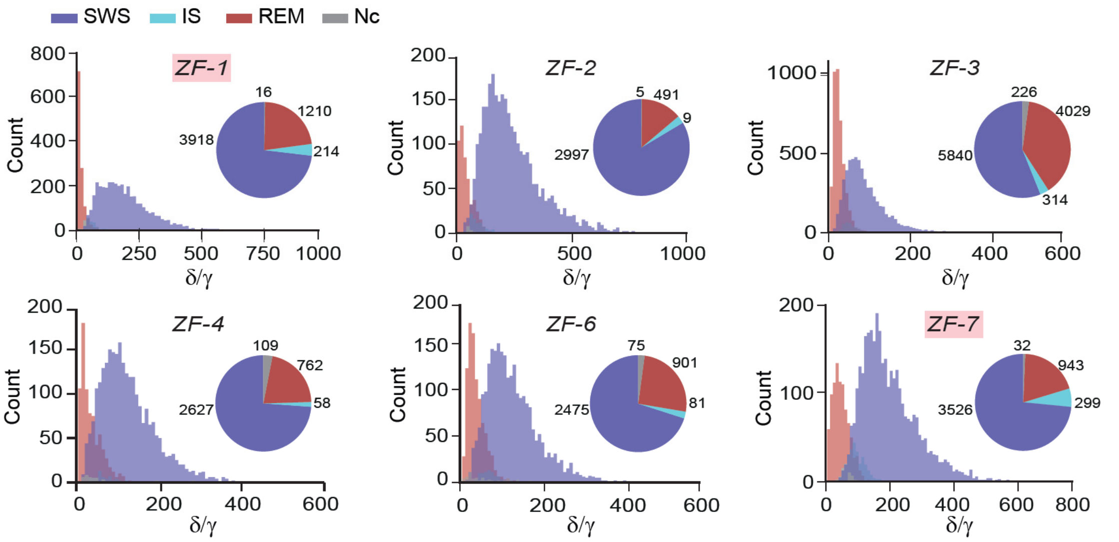


**Supplementary Figure S3: Comparison of sleep stage classification across birds**

Histogram of δ/γ values are grouped according to identified sleep states together with the proportions of sleep stages depicted in a pie chart for all birds. Sub-adult birds are indicated with pink shading of their names. Data from ZF-5 is depicted in Fig. 2.

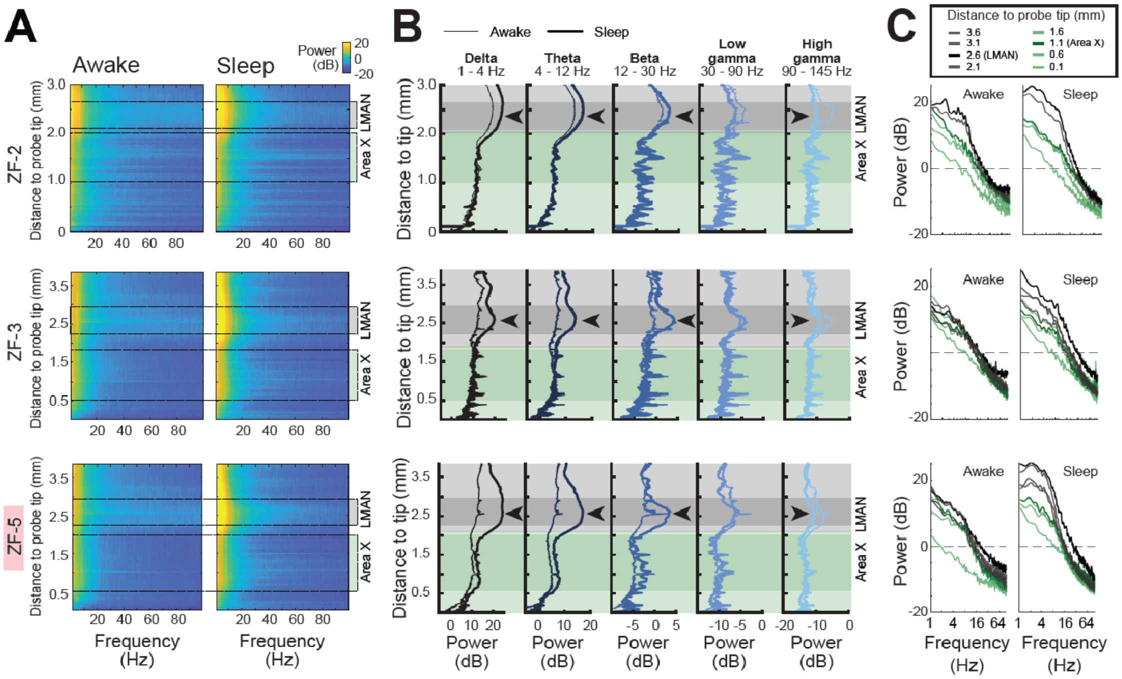


**Supplementary Figure S4: Comparison of spectral features for awake and sleep periods across birds.** Figure conventions same as in Fig. 3 B-D. Subadult ZF-5 is indicated with pink shading of its name.

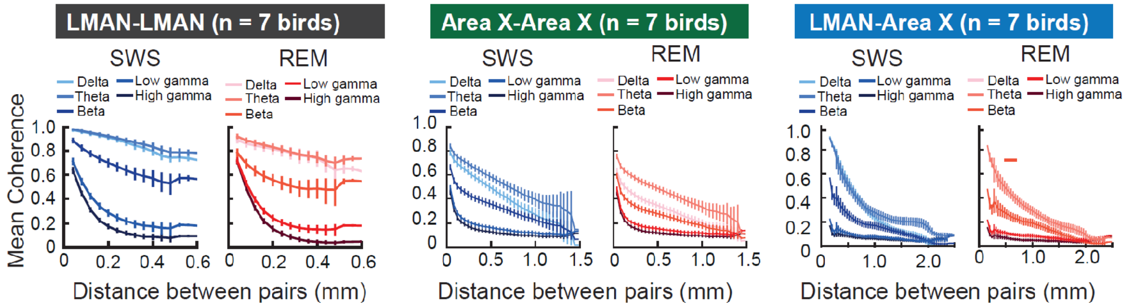


**Supplementary Figure S5: Comparison between REM and SWS coherences for different passbands and as a function of electrode distance.** Panels indicate the change in SWS coherence and REM coherences as a function of inter-electrode distance. Different colored lines indicate different frequency bands. Error bars indicate the 95% confidence interval. From left to right: Coherence analysis for within-LMAN comparisons (left), within-Area X comparisons (middle), and Cross-area LMAN - Area X comparisons (right).

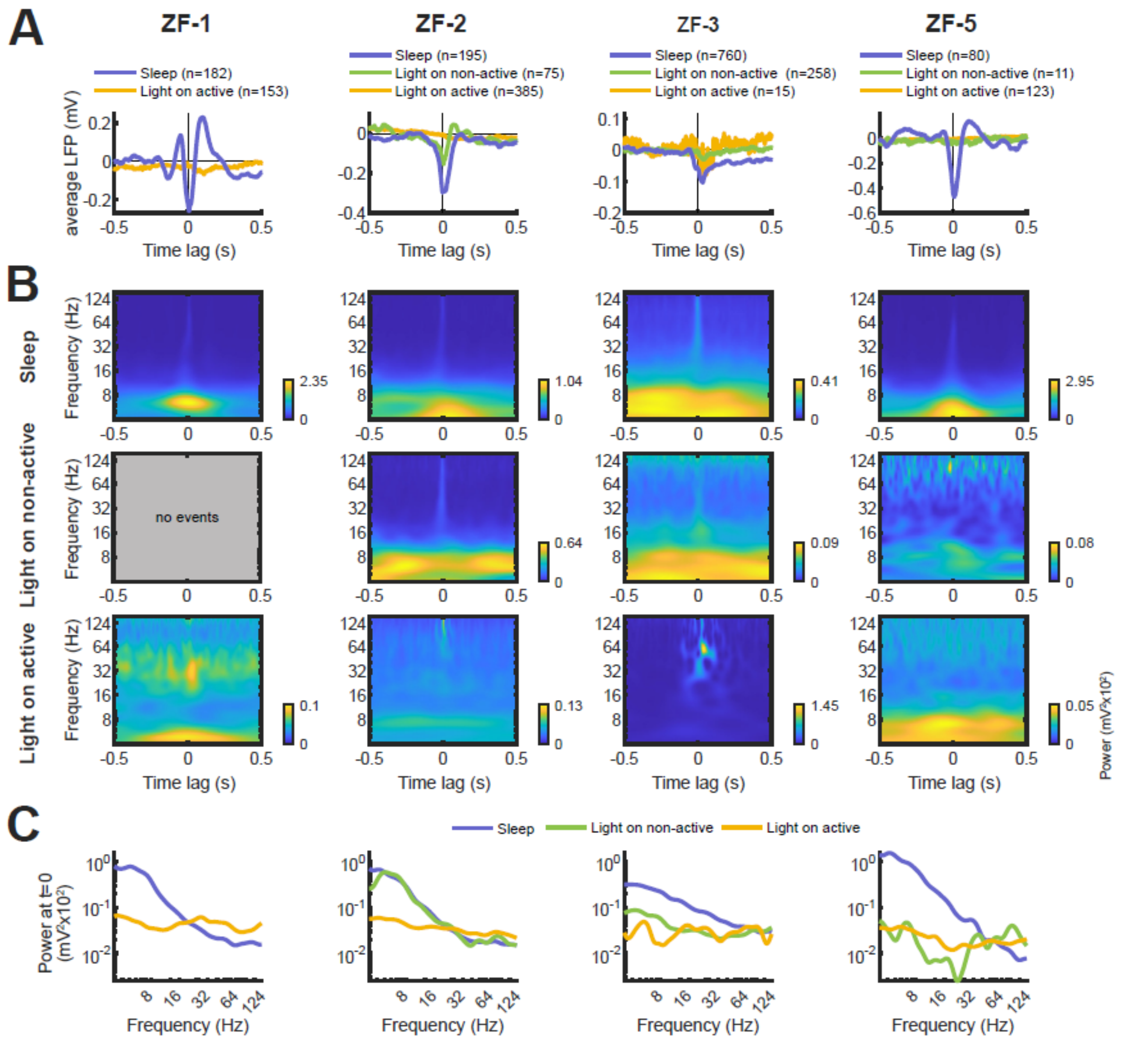


**Supplementary Figure S6: LFP modulation during synchronized bursting in LMAN is state dependent. (A)** Average burst-aligned LFP traces for each behavioral state and animal, centered on the time of synchronized bursting. During sleep (purple), LFPs are strongly modulated, characterized by the prominent negative deflection at t=0. Modulation is variable modulated during light on non-active (green) and absent during light on active periods (yellow). **(B)** Average spectrotemporal power distribution (scalogram) of the burst-aligned LFP across behavioral states, shown separately for each bird. This upregulation is variable across birds during light on non-active (middle row), ranging from a level comparable to sleep in ZF-2, to a reduced level in ZF-3, and nearly absent in ZF-5. No events were detected for ZF-1 for this period. In contrast, during light on active periods (bottom row), no modulation of low-frequency power is observed in any bird. Note that each scalogram is color-scaled individually. **(C)** Power distribution at t=0 take from the scalograms in **(B)** for direct comparison

**Supplementary Table S1: Quantification of sound detections for all recordings**

| | *Light off* | | | *Light on* | | |
|---|---|---|---|---|---|---|
| **Bird** | **Recording length [s]** | **Sound on segments [s]** | **% sound on** | **Recording length [s]** | **Sound on segments [s]** | **% sound on** |
| **ZF1** | 5373 | 105 | 1.95 | 1904 | 1368 | 71.85 |
| **ZF2** | 3855 | 232 | 6.02 | 7595 | 1107 | 14.58 |
| **ZF3** | 10426 | 12 | 0.12 | 3658 | 306 | 8.37 |
| **ZF4** | 3599 | 49 | 1.36 | | | |
| **ZF5** | 3395 | 20 | 0.59 | 2892 | 1405 | 48.58 |
| **ZF6** | 3599 | 41 | 1.14 | | | |
| **ZF7** | 4999 | 86 | 1.72 | | | |

Sound detections statistics that accompany Fig. 3A and Fig. S1.

| | *Light off* | | | | | | *Light on* | | | |
|---|---|---|---|---|---|---|---|---|---|---|
| **Bird** | **Recording length [s]** | **S3 (SWS)** | **S4 (IS)** | **S5 (REM)** | **S99 (NC)** | **S100 (ART)** | **Recording length [s]** | **S0 (REST)** | **S1 (ACT)** | **S100 (ART)** |
| **ZF1** | 5373 | 3918 | 214 | 1210 | 13 | 18 | 1904 | 489 | 1009 | 406 |
| **ZF2** | 3855 | 2990 | 92 | 462 | 4 | 270 | 7595 | 6358 | 1006 | 231 |
| **ZF3** | 10426 | 5840 | 314 | 4029 | 225 | 18 | 3658 | 3006 | 177 | 475 |
| **ZF4** | 3599 | 2627 | 58 | 762 | 109 | 43 | | | | |
| **ZF5** | 3395 | 2427 | 106 | 834 | 6 | 21 | 2892 | 1415 | 1369 | 108 |
| **ZF6** | 3599 | 2475 | 81 | 901 | 72 | 69 | | | | |
| **ZF7** | 4999 | 3526 | 299 | 943 | 31 | 200 | | | | |

Sound statistics for all recordings (Absolute value (s)).

| | *Light off* | | | | | | *Light on* | | | |
|---|---|---|---|---|---|---|---|---|---|---|
| **Bird** | **Recording length [s]** | **S3 (SWS)** | **S4 (IS)** | **S5 (REM)** | **S99 (NC)** | **S100 (ART)** | **Recording length [s]** | **S0 (REST)** | **S1 (ACT)** | **S100 (ART)** |
| **ZF1** | 5373 | 0.73 | 0.04 | 0.23 | 0 | 0 | 1904 | 0.26 | 0.53 | 0.21 |
| **ZF2** | 3855 | 0.78 | 0.02 | 0.12 | 0 | 0.07 | 7595 | 0.84 | 0.13 | 0.03 |
| **ZF3** | 10426 | 0.56 | 0.03 | 0.39 | 0.02 | 0 | 3658 | 0.82 | 0.05 | 0.13 |
| **ZF4** | 3599 | 0.73 | 0.02 | 0.21 | 0.03 | 0.01 | | | | |
| **ZF5** | 3395 | 0.71 | 0.03 | 0.25 | 0 | 0.01 | 2892 | 0.49 | 0.47 | 0.04 |
| **ZF6** | 3599 | 0.69 | 0.02 | 0.25 | 0.02 | 0.02 | | | | |
| **ZF7** | 4999 | 0.71 | 0.06 | 0.19 | 0.01 | 0.04 | | | | |

Sound statistics for all recordings (values are relative to full recording time).

Terminology:

***Light off recordings***

S3 -> SWS
S4 -> IS
S5 -> putative REM, silent rest during light off
S99-> non-classified during sleep segmentation
S100 -> movement artefact (sound amplitude>threshold)

***Light on recordings***

S1 -> active, i.e. vocalizing or moving (sound amplitude>threshold)
S0 -> silent rest (sound amplitude<threshold)
S100 -> LFP artefact (excluded from all analysis)

**Supplementary Table S2: Breakdown of data used in coherence analysis**

| Bird | Total dur (s) | Total REM dur (s) | Total SWS dur (s) | Total REM analyzed (s) | Total SWS analyzed (s) | Total data analyzed (REM+SWS) (%) | Total data analyzed (% of total recording dur) |
|---|---|---|---|---|---|---|---|
| **ZF-1** | 5364 | 1210 | 3918 | 1050 | 3150 | 81,90 | 78,29977629 |
| **ZF-2** | 3855 | 491 | 2997 | 378 | 2646 | 86,70 | 78,44357977 |
| **ZF-3** | 10426 | 4029 | 5840 | 3435 | 3435 | 69,61 | 65,89295991 |
| **ZF-4** | 3599 | 762 | 2627 | 549 | 2196 | 81,00 | 76,27118644 |
| **ZF-5** | 3397 | 835 | 2427 | 729 | 2187 | 89,39 | 85,84044745 |
| **ZF-6** | 3599 | 901 | 2475 | 696 | 2088 | 82,46 | 77,35482078 |
| **ZF-7** | 4999 | 943 | 3526 | 714 | 2856 | 79,88 | 71,41428286 |

**Supplementary Text S1: How would the results be affected if periods of quiet wakefulness were included in the light-off recordings, particularly in segments classified as REM, IS, or SWS?**

Because periods of quiet wakefulness share spectral features with REM, we would expect that awake segments would cluster more closely with REM than with SWS. Importantly, we did not observe any emergence of a distinct fourth cluster that would correspond to wakefulness.

In k-means feature space, adding awake-like points to REM would shift the REM centroid away from its true position. As k-means clustering aims to minimize the summed within-cluster variance, we expect this shift to primarily affect boundaries with IS: if quiet wake lies just outside the REM cluster, REM would broaden and absorb nearby IS points; if quiet wake lies within REM, the cluster would narrow and some borderline REM points would be reassigned to IS, which could in turn draw a few borderline SWS points into IS.

It is important to note that the data form a **continuum in feature space** rather than being separated by sharp boundaries, as illustrated in Figure 2G,H. As a result, some degree of misclassification at the borders is almost inevitable, even without contamination. What contamination would do is to accentuate this tendency by altering the cluster boundary between REM and IS, while leaving SWS relatively stable at the opposite end of the spectrum.

IS appears less prominently in our recordings compared to prior EEG studies. We attribute this to the larger delta range captured by LFP signals in LMAN, which allows periods that might appear as IS in EEG to be more clearly assigned to either SWS or REM (Fig. 3, Fig. S2). Thus, IS in our dataset does not appear artificially inflated; rather, it is more precisely delineated, making REM the most likely recipient of any undetected quiet wake segments.

If our REM sleep periods also included many periods of quiet awakening, we should expect to see that the night time REM coherence and the day time non-active coherence should be similar. As we point out in the discussion, this is not the case: REM and light on non-active coherency values are significantly different across brain regions and passbands (Fig. 4D, see Table 3 for exact values). Therefore, while we have been careful to interpret our results, we are confident that the data that we are analyzing in the dark capture predominately sleep-related activity.

In summary, while some contamination of light-off recordings by quiet awake periods cannot be excluded, our analyses suggest that its impact is minor and does not change the primary conclusions of this study. Misclassifications would occur primarily at cluster borders, with the main effect being a shift at the border between REM and IS. Crucially, the cluster structure appeared intact in all birds, and we did not observe the emergence of a distinct “awake” cluster. We also considered potential effects on coherence results and conclude that any contamination is unlikely to qualitatively alter our findings.

We aim to maximize transparency by providing all raw and preprocessed data, together with the full Matlab and Python code used to generate our figures and analyses in our GIN repository (gin.g-node.org/E4-Leblois/npx_sleep_2024). We hope this will enable readers to fully evaluate our conclusions and facilitate replication.
